# Structural determinants of calcium binding beyond the EF-hand binding site: a study of alpha parvalbumins

**DOI:** 10.1101/2020.09.18.304063

**Authors:** Kalyan Immadisetty, Bin Sun, Peter Kekenes-Huskey

## Abstract

Parvalbumin (PV) is a calcium binding protein expressed in humans, fish and avian species. In these organisms, the calcium (Ca^2+^) affinities of specific PV isoforms can vary by orders of magnitude. Despite the availability of high resolution structural data for many PV isoforms, the structural bases for how such proteins confer widely-varying divalent Ca^2+^ affinities and selectivities against common ions like magnesium (Mg^2+^) has been difficult to rationalize. We therefore conducted molecular simulations of several *α*-pavalbumin (*α*-parvalbumin (*α*PV)) constructs with Ca^2+^ affinities in the micromolar to nanomolar ranges to identify properties of conformations that contribute to their wide-ranging binding constants and selectivities against Mg^2+^. Specifically, we examined a D94S/G98E construct with a reported lower Ca^2+^ affinity (*≈ −*18.2 kcal/mol) relative to the WT (*≈ −*22 kcal/mol), an S55D/E59D variant with enhanced affinity (*≈ −*24 kcal/mol), and a truncated variant of *α*PV with weak affinity (*≈ −*12.6 kcal/mol). We performed molecular dynamics simulations of these constructs and assessed their Ca^2+^ and Mg^2+^ binding properties using scores from molecular mechanics generalized Born approximation (MM/GBSA), ion/oxygen coordination patterns and thermodynamics via mean spherical approximation (MSA) theory, as well as via metrics of protein structure and hydration. Our key findings are that although MM/GBSA and MSA scores successfully rank-ordered the variants according to their previously-published affinities and Mg^2+^ selectivity, importantly, properties of Ca^2+^ loops in CBPs such as coordination, and charge are alone insufficient to rationalize their binding properties. Rather, Ca^2+^ affinity and selectivity against Mg^2+^ are emergent properties stemming from both local effects within the proteins’ ion binding sites as well as non-local contributions from protein folding and solubility. Our findings broaden our understanding of the molecular bases governing *α*PV ion binding that are likely shared by many Ca^2+^ binding proteins.

## 2 Introduction

Parvalbumin (PV) comprise a family of proteins that shape Ca^2+^ signaling in many physiological processes, including neural function, contractility, and immune responses [1]. Given the breadth of PV isoforms found in nature with dramatically different Ca^2+^ and Mg^2+^ affinities [2, 3, 4, 5], these proteins have been popular targets for biochemical and structural characterizations to elucidate the molecular basis of Ca^2+^-affinity [6, 7, 8, 9, 10, 11], or as gene therapies for diseases involving calcium dysregulation [12, 13]. Alpha PV (*α*PV) is a member of the PV family and is expressed in fast twitch muscle fibers, GABA-ergic neurons and the organ of Corti [14]. *α*PV is unique in its extraordinarily high Ca^2+^ affinity relative to other PV isoforms and selectively binds Ca^2+^ over Mg^2+^ by *≈*10 kcal/mol [15, 16]. *α*PV has also been a target for detailed mutagenesis studies [17, 16, 15] to rationalize its Ca^2+^ binding properties including site-directed mutations found to reduce (D94S/G98E *α*PV) or further increase (S55D/E59D *α*PV) [15] the intrinsic Ca^2+^ affinity of *α*PV, as well as a truncated construct (Δ*α*PV) that exhibited dramatically reduced Ca^2+^ binding [18].

Many Ca-binding protein (CBP)s including *α*PV contain helix-loop-helix ‘EF-hand’ metal-binding motifs that have evolved too strongly and selectively bind Ca^2+^. EF hands have a rich evolutionary history that reflects a high degree of sequence homology across protein types, isoforms and species [19]. Ca^2+^ affinity in CBPs such as *α*PV has been shown to be strongly dependent on an EF-hand’s sequence identity, binding site loop length and tertiary structure (reviewed in [20, 21]). Further, selective binding of Ca^2+^ over other mono- and divalent ions, including Mg^2+^, are believed to leverage structural properties of the Ca^2+^ binding loops in EF hands, such as the placement of ion-coordinating polar ligands that exploit the unique size and charge density of Ca^2+^ [21, 22]. Beyond the Ca^2+^ binding loops, myriad structural and dynamic factors tune the affinity of Ca^2+^ binding, including the packing of EF-hand helices, the thermodynamics of exchanging coordinated solvent with metal ion, modulation of secondary structural elements, and structural coupling between adjacent EF-hands [20]. However, the degree to which these factors contribute to the high Ca^2+^ affinities exhibited have not been systematically examined in *α*PV. Insights into these factors could not only inform upon PV proteins’ functions in buffering intracellular Ca^2+^ levels [15], but could extend our understanding of Ca^2+^ sensing in the broad CBP family.

Predicting the energetics of Ca^2+^ and Mg^2+^ binding and structural changes that follow could give important insights into PV proteins, but has remained challenging. Bioinformatics techniques have made strides in identifying binding domains in Ca^2+^ binding proteins based on protein sequences [23, 24, 23]. Other algorithms that account for the geometric configuration of Ca^2+^ chelating ligands to achieve optimal coordination geometries have further improved EF-hand prediction [25] and have helped to rationalize selectivity against other ions like Mg^2+^ [26]. Molecular dynamics (MD) simulations [27] have facilitated detailed analyses of Ca^2+^ binding in intact proteins, using force fields fit to quantum chemical calculations of metal oxygen interactions [28]. More recent applications have focused on the refinement of Ca^2+^ force field parameters [29, 30], energetics of Ca^2+^ binding in EF-hand containing proteins [31, 32, 33], and effects of Ca^2+^ binding on protein function [34, 35, 36].

Elkins et al. [17] presented one of the first computational studies of PV to leverage molecular simulations. In this computational mutagenesis study, the residue in the ninth position (i.e., E59) of the silver hake (SHPV-B) PV proteins’ first EF hand was mutated to arginine/serine/cysteine in order to demonstrate that the pairing of amino acids comprising the first and second EF hands modulated the proteins’ apparent Ca^2+^ affinity. Cates et al. extended these studies to probe the relationship between coordinating ligands of the EF hand loops and the overall flexibility and plasticity of pike PV via MD simulations [37] to show that the EF hands inherent rigidity is preventing bidentate ligation when the residue is mutated to aspartate. More recently, Kucharski et al. utilized MD simulations and a statistical mechanical model, mean spherical approximation (MSA), to rationalize *β*-parvalbumin (*β*PV)s’ thermodynamic preference for Ca^2+^ over Mg^2+^ and concluded that EF hand rigidity disfavors binding of the smaller Mg^2+^ ion and thereby contributes to its selectivity for Ca^2+^ [38]. Other studies have used methods to model the structure of EF hand motifs in PVs [39] and probe the folding/unfolding in *β*PV. To date, no such studies have been conducted for the high affinity *α*PV variants.

In this study, we apply MD simulation and MSA approaches developed in [38] to rationalize trends in Ca^2+^ binding among *α*PV variants and general constructs. We considered variants that exhibit strongly differing Ca^2+^ affinities, including two *α*PV variants (S55D/E59D *α*PV and D94S/G98E *α*PV) with mutations within their Ca^2+^ binding loops, as well as a truncated variant, Δ*α*PV, which lacks two helices (A,B). Our studies indicate that while features intrinsic to the EF hands strongly contribute to Ca^2+^ binding and selectivity in *α*PV, allosteric changes accompany the observed affinity changes.

## 3 Results

Our primary goal in this study was to determine whether molecular simulations could provide mechanistic insights into the Ca^2+^ binding affinities and selectivities versus Mg^2+^ in *α*PV. Towards this end, we performed nearly *≈*36 *µ*s of unbiased MD simulations for the Ca^2+^-bound, Mg^2+^-bound and apo states of *α*PV, as well as several constructs (S55D/E59D *α*PV, D94S/G98E *α*PV, and Δ*α*PV), for which structures have been reported in the literature [15, 16, 18, 40]. Since atomistic resolution structural data have only been determined for a subset of these variants, we first compare our predicted structures with those available in the Protein Databank. We then test whether scores estimated via MM/GBSA and MSA theory can rank the *α*PV structures in a manner consistent with their experimentally-determined Ca^2+^ affinities. We next decompose these scores into contributions from the EF-hand Ca^2+^ coordination sites versus structural and energetic changes elsewhere in the protein.

### 3.1 Comparison of simulated *α*PV structures

We first verify that the MD predicted structures resemble those deposited in the Protein Databank. In Fig. 1A,B we demonstrate that the predicted apo (blue) and holo (red) WT structures are generally consistent with their respective experimental reference structures (PDB:2JWW [41] and PDB:1RWY [40] (green)). The similarity is also evidenced by the small root mean squared deviations (RMSD) values of roughly 2.0 and 1.0 Å for the apo and holo states, respectively (Fig. S1). We note similar secondary structure content in the *α*PV variants relative to the wild-type (WT), including the *β*-sheets linking the CD and EF domain loops as well as the high *α*-helical content (see sub-panel in Fig. 1A).

**Figure 1:**
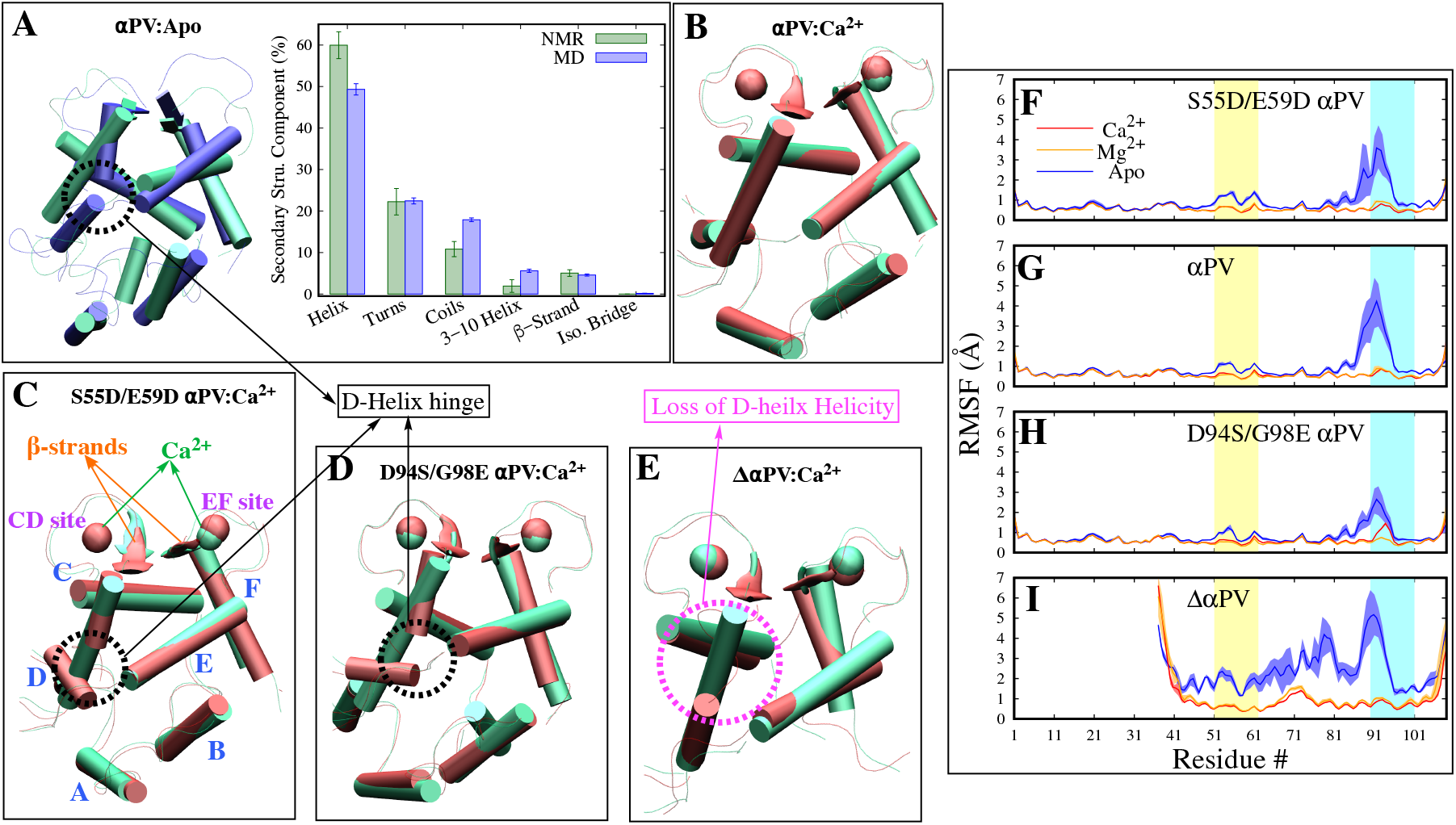
**A-E** Comparison of experimental (NMR/X-ray) and molecular dynamics (MD) predicted *α*PV structures. All experimental structural snapshots are shown in green and MD snapshots are shown in either blue (apo) or red (holo). **A-E** Representative snapshots of NMR and MD *α*PV apo systems were superposed and their secondary structural components are shown. Each MD data point is the average and standard deviation calculated from the means of three trials and the NMR data point is the average and standard deviation of all NMR modes. **F-I** RMSF versus residue number. All apo are colored blue and Ca^2+^ bound systems red and Mg^2+^ bound orange. Average of three trials of each system is shown; standard deviation is shown as shaded curves. Yellow and cyan shaded regions are CD and EF site loop regions. Residues spanning the different parts of the protein are mapped as follows: Helix A (S1-A17), helix B (D25-G34), helix C (A40-L50), helix D (E59-F70), CD site loop (D51-I58), helix E (S78-D90), helix F (V99-E108), and EF loop (K91-G98).

We next evaluated in Fig. 1 the per residue conformational dynamics of the proteins via root mean squared fluctuations (RMSF) to identify potential differences among the *α*PV variants that could correlate with ion affinity. The RMSFs indicate that bound Ca^2+^ in the holo states expectedly suppress fluctuations in all variants relative to the apo states. Similar to observations in other closely related Ca-binding proteins (CBPs) including troponin C (TnC) [36, 42] and *β*PV [38], the reduction in RMSF upon ion binding is largely restricted to the EF hand loops. Here, the bound ions stabilize the CD (residues L50 to E60) and EF (residues L85 to G95) loop regions (Fig. 1). The most significant RMSF changes occur in the EF site loop, as the average maximal peak height in the EF site loop in the apo forms is *≈*5Å compared to *≈*1.5Å for the holo. Further, we noted a modest correlation between the apo-state RMSF values and Ca^2+^ affinity in the CD and EF sites, as variants with stronger Ca^2+^ yielded larger values relative to those with weaker affinities. We did not observe any statistical difference between the Ca^2+^ and Mg^2+^ bound forms (Fig. 1F-I). Overall, these data indicate modest correlation between the extent by which RMSFs are reduced upon Ca^2+^ binding and experimentally-measured binding affinities. However, as a whole, RMSF differences were small across all variants irrespective of the type of bound ion.

### 3.2 Scoring *α*PV variants’ binding affinites

Our simulations suggest that ion binding reduces the RMS fluctuations intrinsic to each *α*PV apo state. Here we used the end-point thermodynamics method, MM/GBSA, to determine energetic bases for these changes that may contribute to high Ca^2+^ and Mg^2+^ affinity. We first tested whether energies predicted from the simulations via the MM/GBSA method could reliably rank-order the variants by their experimentally-determined affinities. In Table S1 we provide full MM/GBSA energies representing the internal, van der waals, electrostatic, and polar- and non-polar solvation interaction terms as free energies 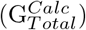 or 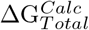 between the *α*PV/Ca^2+^ complex (holo) and apo structure. Note our 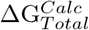 does not include contributions for the solvation energy of the free ions, as these terms are the same for each *α*PV variant. To compare variants, we report ΔΔ*G ≡* Δ*G*_*i*_ *−* Δ*G*_*j*_, which cancels out contributions from the free Ca^2+^s’ solvation energy. The estimated MM/GBSA scores 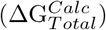 unfortunately failed to correctly rank the proteins, which we attributed to the MM/GBSA estimates for the ion/protein interactions, as explained below.

The Generalized Born Surface Area approach used for the MM/GBSA scoring was developed as a fast alternative to computing electrostatic energies from Poisson-Boltzmann (PB) theory [43]. However, standard PB theory suffers from inaccuracies when applied to highly charged systems [44], such as the Ca^2+^-bound EF-hand loops we considered here. Therefore, we determined if omitting the Ca^2+^/protein binding interaction terms would provide a more accurate ranking, by assessing the ‘reorganization score’ of the protein upon binding an ion. We define this reorganization score as the difference in MM/GBSA score of the isolated protein in its holo state versus that of its apo form. Indeed, the reorganization energies yielded 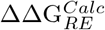 values of *−*19.94 *±* 0.82, 0.0 *±* 0.84, and 30.04 *±* 0.84 kcal/mol for S55D/E59D *α*PV, *α*PV, and D94S/G98E *α*PV, respectively (Table 1), which suggest that forming the holo state in S55D/E59D *α*PV incurs a smaller thermodynamic penalty than the WT, while Δ*α*PV incurs an even greater cost. The trend of reduced reorganization score correlates with the experimental affinity estimates of (ΔΔG^*Expt*^) of *−*2.0 *±* 0.22, 0.0 *±* 0.08 and 3.8 *±* 0.22 kcal/mol (Table 1). We further computed a 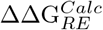 of *−*0.26 *±* 0.76 kcal/mol for the Δ*α*PV. However, the apo Δ*α*PV structures we simulated remained partially folded and thus did not represent the unfolded state suggested by circular dichroism experiments [18], thus its 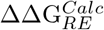 was not meaningful. For this reason, in subsequent analyses of Δ*α*PV we only consider the score decomposition of its holo state (i.e., 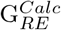 in Table 1). We observed roughly similar trends for Mg^2+^, though D94S/G98E *α*PV binds the ion with a modestly higher score relative to the WT. Importantly, we note that 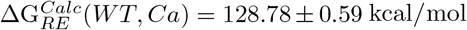 versus 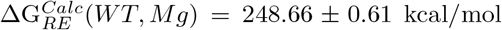. These scores suggest that binding Mg^2+^ manifests in an even greater reorganization penalty relative to Ca^2+^ that would disfavor the binding of magnesium.

**Table 1:**
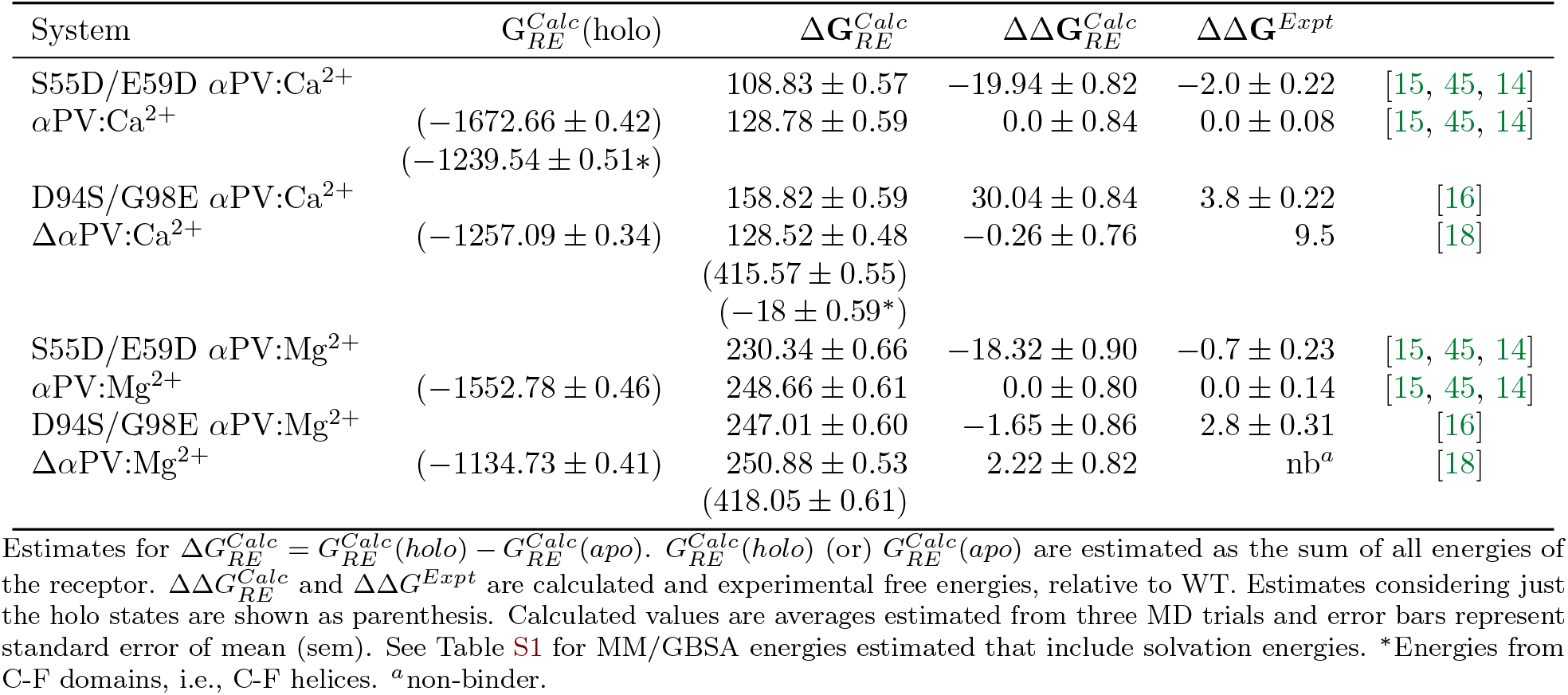
Estimation of MM/GBSA reorganization energies.

| System | $G_{RE}^{Calc}(\text{holo})$ | $\Delta G_{RE}^{Calc}$ | $\Delta\Delta G_{RE}^{Calc}$ | $\Delta\Delta G^{Expt}$ | |
| --- | --- | --- | --- | --- | --- |
| S55D/E59D αPV:Ca<sup>2+</sup> | | $108.83 \pm 0.57$ | $-19.94 \pm 0.82$ | $-2.0 \pm 0.22$ | [15, 45, 14] |
| αPV:Ca<sup>2+</sup> | $(-1672.66 \pm 0.42)$<br>$(-1239.54 \pm 0.51^*)$ | $128.78 \pm 0.59$ | $0.0 \pm 0.84$ | $0.0 \pm 0.08$ | [15, 45, 14] |
| D94S/G98E αPV:Ca<sup>2+</sup> | | $158.82 \pm 0.59$ | $30.04 \pm 0.84$ | $3.8 \pm 0.22$ | [16] |
| $\Delta\alpha$PV:Ca<sup>2+</sup> | $(-1257.09 \pm 0.34)$ | $128.52 \pm 0.48$<br>$(415.57 \pm 0.55)$<br>$(-18 \pm 0.59^*)$ | $-0.26 \pm 0.76$ | 9.5 | [18] |
| S55D/E59D αPV:Mg<sup>2+</sup> | | $230.34 \pm 0.66$ | $-18.32 \pm 0.90$ | $-0.7 \pm 0.23$ | [15, 45, 14] |
| αPV:Mg<sup>2+</sup> | $(-1552.78 \pm 0.46)$ | $248.66 \pm 0.61$ | $0.0 \pm 0.80$ | $0.0 \pm 0.14$ | [15, 45, 14] |
| D94S/G98E αPV:Mg<sup>2+</sup> | | $247.01 \pm 0.60$ | $-1.65 \pm 0.86$ | $2.8 \pm 0.31$ | [16] |
| $\Delta\alpha$PV:Mg<sup>2+</sup> | $(-1134.73 \pm 0.41)$ | $250.88 \pm 0.53$<br>$(418.05 \pm 0.61)$ | $2.22 \pm 0.82$ | nb<sup>a</sup> | [18] |

### 3.3 Ca^2+^ affinity determinants within the EF hands

Since both site-directed mutations in S55D/E59D *α*PV and D94S/G98E *α*PV reside within the EF hands, it was expected that changes in binding free energy could be explained by differences in the binding loop conformations that coordinate Ca^2+^. We therefore measured the radial distribution of protein oxygens around bound ions in the CD site to assess the loops’ coordination of Mg^2+^ and Ca^2+^ for the S55D/E59D *α*PV(red), *α*PV(green), D94S/G98E *α*PV(blue) and Δ*α*PV(cyan) proteins in Fig. 2. We observed that the radial distribution maxima for the *α*PV variants occurred at smaller Ca^2+^/oxygen distances (*≈*2.2 Å) relative to that of the WT (*≈*2.4 Å) (Fig. 2A). However, the cumulative oxygen density (Fig. 2C) obtained by integrating the radial distributions with respect to distance (Fig. 2A) indicate that within 3 Å, the WT has a near optimal coordination number of eight oxygens, while the D94S/G98E *α*PV and Δ*α*PV cumulative densities reflect seven bound oxygens. Curiously, the high affinity S55D/E59D *α*PV variant bound Ca^2+^ with just six oxygens (Fig. 2C). We observed similar trends in the EF site, namely that the corresponding cumulative distributions indicate D94S/G98E *α*PV coordinates seven oxygens within 3.0 Å of Ca^2+^, whereas for all other systems only six are coordinated (Fig. 2D). Assessments for Mg^2+^ are summarized in Section S8.4. These lack of consensus in these data suggest that the protein coordination numbers alone do not strongly correlate with affinity. Similar analyses enumerating the waters complexed within the EF hands were also inconclusive.

**Figure 2:**
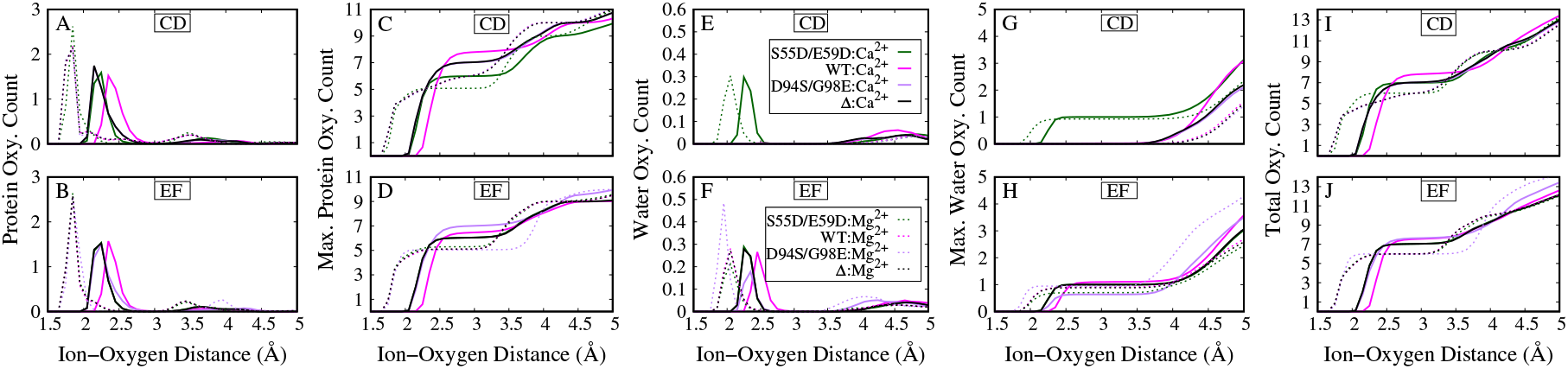
Protein and water co-ordination with the ions in the CD and EF sites. **A, B** Radial distribution of protein oxygens in the CD (E) and EF (F) sites. **C, D** Maximal amino acid oxygen count in CD (G) and EF (H) sites. **E, F** Radial distribution of water oxygens in the CD (A) and EF (B) sites. **G, H** Maximal water oxygen count in CD (C) and EF (D) sites. **I, J** Total oxygen count, i.e., the sum of water and amino acid oxygen count in CD (C) and EF (D) sites. All Ca^2+^ bound systems are shown as full lines and the Mg^2+^ bound ones as broken lines. S55D/E59D *α*PV, *α*PV, D94S/G98E *α*PV and Δ*α*PV are colored in green, magenta, purple and black respectively.

The ΔΔ*G* values from MM/GBSA correctly rank the *α*PV proteins by their reorganization energies but only did so by neglecting Ca^2+^/protein interactions as explained earlier. We therefore used an alternative method, Mean spherical approximation (MSA) theory, to estimate the thermodynamics of binding Ca^2+^ and other ionic species to the EF hands in a manner that accounts for the highly-charged and finite-sized atoms in the binding site. Specifically, we use MSA to relate the MD-predicted ion coordination data (numbers of coordinating oxygens and binding site volumes) to the chemical potentials for partitioning solvent ions (potassium (K^+^), Mg^2+^, Ca^2+^ and chloride (Cl^*−*^)) into the EF and CD binding domains. We present in Fig. 3 MSA-predicted chemical potentials for cations in a 150 mM KCl, 1 mM MgCl_2_ and 2 *µ*M CaCl_2_ solution that emulate intracellular ionic strengths. The chemical potentials are presented for several binding site volumes to reflect the range of maxima in the oxygen radial distributions we computed (0.4 to 0.65 nm^3^ to represent radii of 0.25 to 0.36 nm) as well as coordination by five to eight oxygens. Additionally, we varied the average charge of the coordinating oxygens to reflect the contributions of backbone oxygens (−0.57e), serine hydroxyl oxygens (−0.65e), glutamic or aspartic acid oxygens (−0.82), and water oxygens (−0.83e) to the average effective charge of oxygen in the binding domain. Our MD simulation data indicate an effective oxygen charge (*q*eff) of the CD site for the S55D/E59D *α*PV variant of −0.77e, which is more negative than the CD site for the other variants. This difference reflects a greater proportion of oxygens from acidic side chains in the CD site of S55D/E59D *α*PV(while the EF-hands for all cases have *q*eff = −0.75e (see Table S6)).

**Figure 3:**
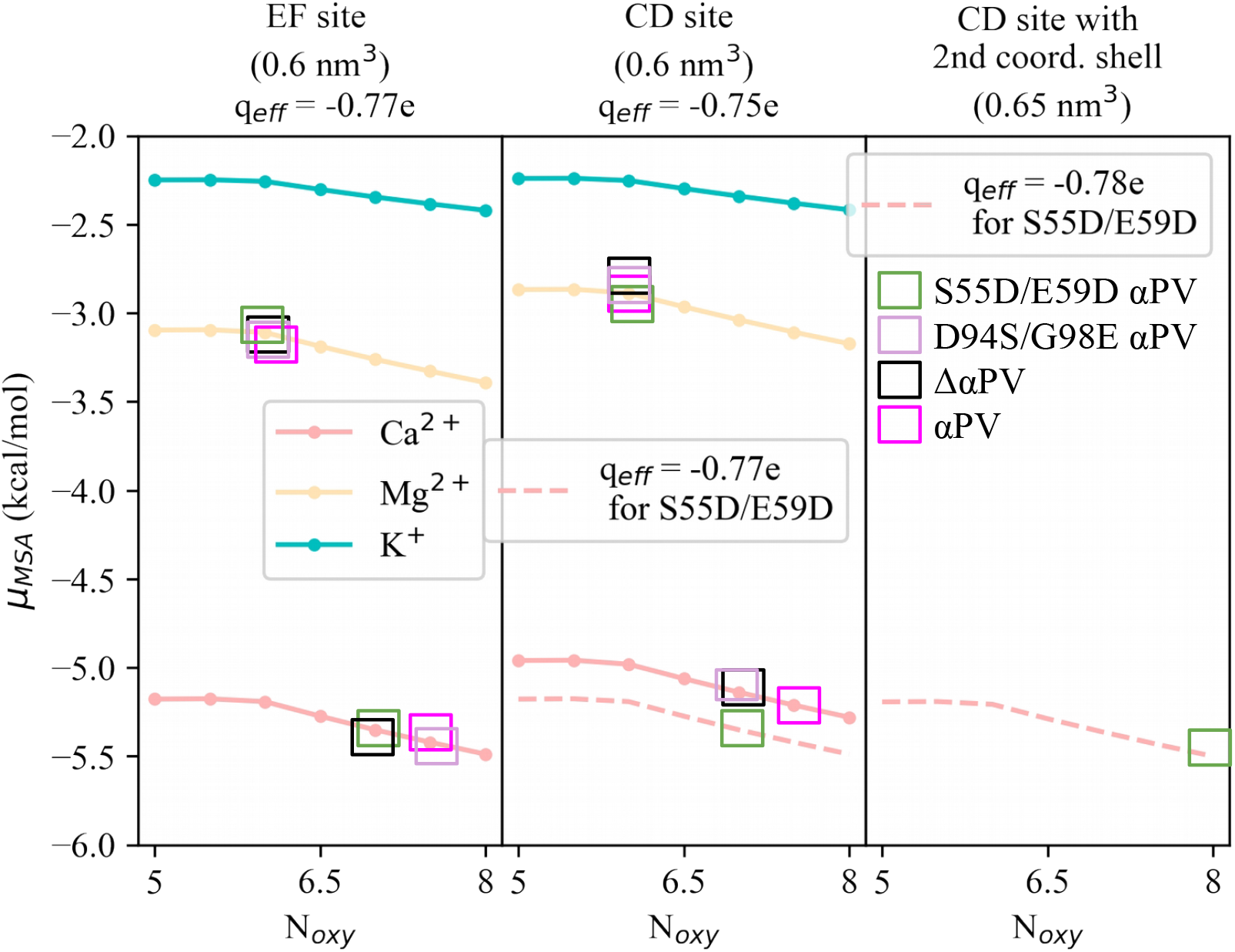
MSA free energies for cations in the *α*PV binding site with bath concentrations [KCl] = 150 mM, [MgCl_2_] = 1 mM and [CaCl_2_] = 1 *µ*M under varying numbers of coordinating oxygens (*N*_*oxy*_) from 5.0 to 8.5. The remaining parameters are the same as our previous MSA calculations in [38]. The colored boxes indicate the *α*PV systems we simulated (including water oxygens).

Our results in Fig. 3 consistently demonstrate that the chemical potentials for Ca^2+^ (red) are more favorable than Mg^2+^ (blue) by approximately 2 kcal/mol over all binding site radii we considered. This difference in large part reflects the higher desolvation energy of Mg^2+^ relative to Ca^2+^ (−1931.4 versus 1608.3 kJ/mol [46]) that offsets the entropic advantage of partitioning the smaller ion into the binding site. This result is consistent with experimental data indicating that Ca^2+^ affinities are *≈*10 kcal/mol (see Tables S1, 1) more favorable than Mg^2+^. Further, binding site configurations with a more prominent negative charge (−0.77e), e.g. for S55D/E59D *α*PV CD site, presented modestly more favorable chemical potentials for divalent cations of roughly 0.5 kcal/mol over all other variants (−0.75e). In other words, the more negative CD binding site of S55D/E59D *α*PV provides thermodynamic stability over the other site configurations and is consistent with the *≈*2 kcal/mol (see Table S1) energy difference reported via experiment. Further, the model indicates that the chemical potential is modestly insensitive to the number of bound oxygens (8 is favored over 5 by less than 1kcal/mol). Since MSA is a coarse approximation of ion binding thermodynamics, it is unclear whether *<* 1 kcal/mol difference in chemical potentials is significantly different for *α*PV. Nonetheless, by accounting for finite size effects and competition with other solvated electrolytes for the proteins’ binding domains via MSA theory, the proteins’ Ca^2+^ selectivity over Mg^2+^and modest differences in the predicted Ca^2+^ chemical potentials can be rationalized. In contrast, the MM/GBSA scores summarized in Section S8.4 suggest that the 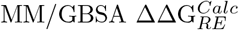 reorganization scores for the CD site, but not the EF site, correlated with the variants’ Ca^2+^ affinities (Table S2).

### 3.4 Ion affinity determinants beyond the EF hands

We have so far demonstrated that MD-predicted differences in the EF-hand coordination and associated reorganization energies among the *α*PV variants loosely correlate with experimental measurements. Nevertheless, we found that point mutations also impart changes in the overall structure of the protein (Fig. 1B-D), as well as its dynamics (Fig. 1F-H). We therefore investigated properties of the entire protein that could further contribute to the range of affinities reported in *α*PV variants, including desolvation, helix bundle ‘packing’, and secondary structure.

Graberek et al. [21] speculated that waters coordinated to solvated Ca^2+^ are released into the bulk medium as the ion binds proteins. These liberated waters increase the configurational entropy of the bulk medium and thereby lower the free energy of the system. In this vein, we investigated analogous changes in the solvation of the *α*PV variants that occur during Ca^2+^ binding. In Fig. 4C, we report the number of waters within 4.0 Å of each protein configuration as an indicator of solvation. Across all systems, we observe at least a reduction (green bars) of twenty bound waters in the ion-bound holo states relative to the apo states. Moreover, there is a significantly greater loss of solvent for the intact proteins that strongly bind Ca^2+^ (*∼*54, *∼*56 for S55D/E59D *α*PV and *α*PV) relative to D94S/G98E *α*PV (*∼*26). Similar trends were observed in Mg^2+^ bound systems, although, no significant differences were found relative to the Ca^2+^ bound systems. These trends in water loss are similar to those observed for the proteins’ radii of gyration (Fig. 4A). Δ*α*PV is an exception to this trend in that at least 50 water molecules were liberated; in all likelihood, this number would be far greater if the unfolded ion-free state were considered. The changes in water solvation appear to arise from a reduction in the proteins’ solvent accessible surface area (SASA) upon ion binding (Fig. S3). Overall, we observe a consistent trend of increased dehydration upon ion binding that correlates with the intact proteins’ increasing Ca^2+^ affinity.

**Figure 4:**
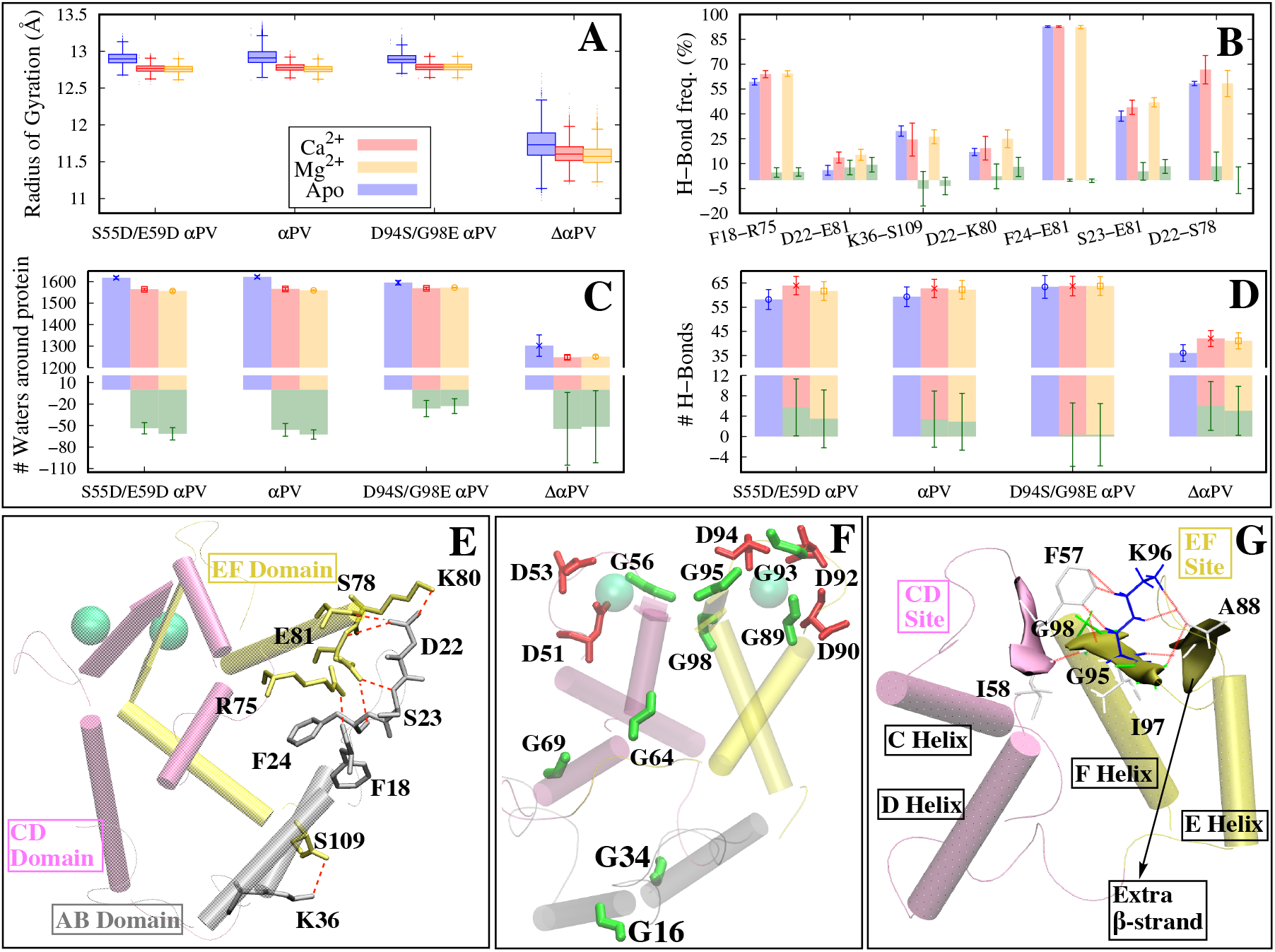
**A** Comparison of radii of gyration (R_*g*_) of all systems. All three trails of a system are plotted as a single box plot. Boxes represent the 25th to 75th quartile (i.e., the interquartile range where 50% of data is concentrated), bottom and top whiskers are minima and maxima, and the horizontal line in the boxes represent the median, and the points above and below whiskers are outliers. **B, E** Hydrogen bonds that are formed between the AB domain (colored silver) and remainder of the protein in the holo *α*PV. CD and EF domains are colored mauve and yellow, respectively. **C** Waters within 4 Å of protein. Each data point represents the mean of three trials and error bar represent standard deviation. Green bars are the difference between holo and apo systems (ΔW). **D** Total number of hydrogen bonds in the last 200ns of simulation. Error bars are standard deviations estimated from three MD trials. Green bars are the difference between holo and apo systems (ΔHB). **F** Glycines (green) and aspartic acids (red) in *α*PV. **G** Extra *β*-strand formed in the EF loop (colored yellow) in the apo Δ*α*PV. Extra *β*-*β* hydrogen bonds that are formed in the EF loop region are colored red and the corresponding residues are shown as sticks. CD and EF domains are colored pink and yellow respectively.

In the previous section, we identified correlated differences in desolvation and SASA upon ion binding. We speculated that these differences could be attributed to isoform-specific configurations of the helices composing the proteins globular domains. We therefore report in Fig. 1B-E the relative positions of helices composing the proteins’ hydrophobic core in order to highlight conformational changes among the variants as they bind Ca^2+^. These changes culminated in more compact proteins during ion binding as evidenced by the reduction of their radii of gyration (Fig. 4). We anticipated these structural changes reorganize the interfaces between bundle helices, and entail reorganization of the hydrogen bonding network within and between these helices. Along these lines, we demonstrate that the number of hydrogen bonds tend to be greater in the holo state relative to apo (green bars in Fig. 4D). However, only for S55D/E59D *α*PV was the change significantly different from zero among the intact proteins. Hence it was apparent that more side chain hydrogen bonds were formed than broken during ion binding, but we did not observe consistent patterns that strongly differentiated weak from strong Ca^2+^ binders. Further, although the average change in hydrogen bond appears to be lower for the Mg^2+^ bound systems, the difference between the two ions is not significant. A summary of specific hydrogen bonding interactions involving side chains is provided in Section S8.4. Overall, we show that upon ion binding increased hydrogen bonding resulting in an overall decrease in the reorganization scores as shown in Table 1.

In Fig. 4D we note that the total number of hydrogen bonds in Δ*α*PV is significantly smaller (*≈*35) compared to the intact systems (*≈*60) as would be expected from loss of its AB helices. Interestingly, the change in hydrogen bonds we report for Δ*α*PV is significantly greater than zero (*≈*6), which suggests Ca^2+^ plays an important role in increasing the connectivity of the hydrogen bond network within the CDEF helices. We expect that the actual change in the number of hydrogen bonds would be far greater if we had assumed an unfolded conformation for the apo state. From this set we identified seven hydrogen bonds spanning the AB/CDEF interface that underwent significant changes in bonding frequency during ion binding (see Fig. 4B). Bonds that exhibited greater contact frequency upon Ca^2+^ binding include F18/R75, D22/E81, D22/K80, D22/S78, S23/E81, and F24/E81, while K36/S109 had fewer contacts. Though nearly all of these reflected increases in bonding frequency in the holo relative to apo states, those changes were generally minor. In other words, the hydrogen bonding network was modestly ‘tightened’ relative to its apo configuration, as opposed to undergoing dramatic changes in the hydrogen bond network. These structural changes culminated in a net gain of *≈*18 kcal/mol upon deleting the AB domain from *α*PV (Table 1 and Fig. S4), as assessed using scores for residues K37/M37 to E108 in *α*PV. Hence, it is apparent that there are hydrogen bonds between AB/CDEF helices in the WT *α*PV that are favorable and undoubtedly help to stabilize the folded state. Some of these appear to have higher bonding frequencies, which may imply the AB helices help to stabilize the hydrogen bonds and other nearby residues in the holo over apo states.

Given the general increase in hydrogen bonding upon complexing metals we reported in the last section, we next assessed changes in *α*-helix and *β*-sheet secondary structure and their corresponding energies. *α*-helices represent the most abundant secondary structure in *α*PV (*≈*45-55%) (Fig. 1A and Fig. 5A). Our MD simulations indicate that binding ions increases the *α*-helical content of all *α*PV variants (*≈*3-6%), with Ca^2+^ to a greater extent than Mg^2+^ (*≈*1-4%) (Fig. 5A). Overall, the increase in *α*-helicity is greater for Ca^2+^ relative to Mg^2+^, while the magnitude of helicity changes is inversely correlated with affinity among the intact variants (5% for D94S/G98E *α*PV, 4% for WT, 3% for S55D/E59D *α*PV).

**Figure 5:**
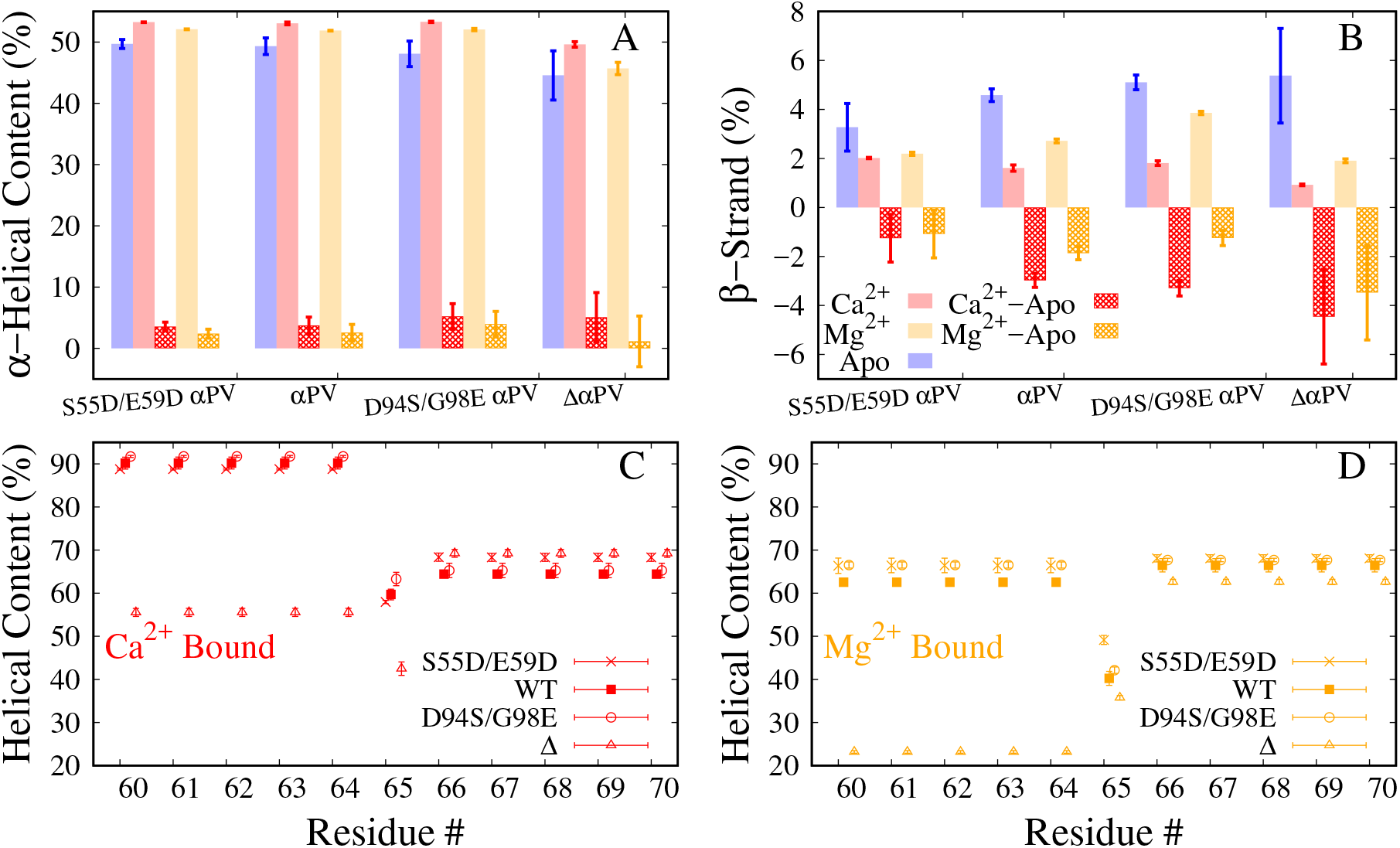
Secondary structural elements of all systems compared. **A-B** *α*-helicity (A) and *β*-strand (extended configuration according to VMD) (B) of the entire protein over the entire simulation time was calculated. Each data point is the average and standard deviation of three trials of each system (taken from means of each trial). The difference between the holo state and apo state is also plotted as grid bars. Only C_*α*_ atoms are considered for this calculation. **C-D** *α*-helix of the D-helix residues for the Ca^2+^ (C) and Mg^2+^ (D) bound systems are compared.

In Fig. S6 we show that most of the changes in helicity during ion binding arise in helix E (residues A79 to G89), and to a lesser extent in helix D (residues E60 to F70). These data indicate *α*-helical character increases in the holo relative to the apo state and those changes are adjacent to the loop residues that directly coordinate Ca^2+^. For instance, the N-terminal residues of helix D (E60-S65) undergo an increase from under 50% helicity to approximately 60 and 90% for Mg^2+^ and Ca^2+^, respectively (Fig. 5C, D) and (Fig. S6) versus more modest changes within S65-F70. The low helicity of the apo state residues E60-G64 (by *≈*20-25%) in the D-helix coincides with the glycine at position 64 (shown in Fig. 4F). Similarly, the N-terminal region of helix E (A79-T84) exhibits *≈*90% or greater *α*-helicity, compared to the C-terminal end that approaches less than 60% at residue A87. These data also suggest that the binding of Ca^2+^ and Mg^2+^ maximize the helical content to nearly 100%. Mg^2+^ reduces the helicity of residues E60-S65 in D-helix at least by 20% in all variants (Fig. 5D). Helices D and E exhibited increased helicity upon ion binding, which in turn yielded favorable 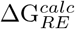 scores that correlated with experimental trends in Ca^2+^ affinity (see Fig. S4 and Table S4).

We also demonstrate that truncating the AB domain from *α*PV reduces the overall helical content, particularly within the D helix. Specifically, as with the intact variants, the greatest impact on the helicity of the D-helix was restricted to the N-terminal end (E60-S65), *∼*55% versus nearly 90% for the intact variant (Fig. 5C). The corresponding energies from MM/GBSA indicated that the D helix was more thermodynamically favored in the WT holo *α*PV relative to Δ*α*PV (Ca^2+^ bound *α*PV and Δ*α*PV are *≈ −*149 and *≈ −*137 kcal/mol respectively for helix D from Table S4). For the E helix, no differences were not apparent (Table S5). These data indicate that the AB domains maintain a high degree of *α*-helicity in the D helix upon ion binding, which likely contributes to high affinity Ca^2+^ binding.

The two *β*-strands in *α*PV couple the two ion binding domains through the formation of anti-parallel *β*-sheet (Fig. 1A). This secondary structure element is common across EF-hand containing proteins and are implicated in shaping Ca^2+^ binding affinity [47, 20]. We observed in *α*PV proteins that the binding of Ca^2+^ or Mg^2+^ both reduce the *β*-strand content in the *α*PV systems by *≈*1-5%, with generally more significant reductions in *β* character for Ca^2+^ binding relative to Mg^2+^(Fig. 5B). This reduction in *β*-strand content likely occurs as a result of F57 (Table S7) and K96 (Table S8) backbone oxygens shifting from the *β*-sheet in the apo state to binding Ca^2+^ in the holo state. *β*-strand content is reduced in residues F57-E59 and K96-G98 most significantly by *≈*50-60% for Ca^2+^ versus *≈*20-40% for Mg^2+^ (Fig. S7). Additionally, we note that the reductions in *β* sheet character are inversely correlated with the Ca^2+^ binding affinity of the *α*PV variants, although the effect size by percentage is relatively small (*≈ −*1.0% for S55D/E59D *α*PV relative to *≈ −*3% for D94S/G98E *α*PV). We had anticipated that the more extensive unfolding of the *β*-strands in weaker versus stronger variants would yield thermodynamically unfavored score changes. As expected, we found that the associated changes in 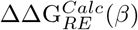 for these residues correlated with affinity (*−*17 kcal/mol for S55D/E59D *α*PV versus 28 kcal/mol for D94S/G98E *α*PV in Table S3) Hence, we observe potentially meaningful trends in the extents of *β*-sheet unfolding during ion binding that correlate with the variants’ respective affinities.

## 4 Discussion

*α*PV is among the highest affinity binders of Ca^2+^ in the PV family. The two site-directed mutants evaluated in our study include S55D/E59D *α*PV and D94S/G98E *α*PV, which bind Ca^2+^ with stronger and weaker affinities compared to the WT *α*PV, respectively. Both variants reflect substitutions in the Ca^2+^ binding EF-hand loops, which would generally be expected to directly impact Ca^2+^ affinity by changing the ions’ coordination. A truncated *α*PV variant (Δ*α*PV) was also considered to investigate the role of the AB helices indirectly influencing the binding of Ca^2+^. Our primary goals of this study were to 1) test the hypothesis that Ca^2+^ affinity is significantly modulated by structural features beyond the ion-coordinating EF hand loops, 2) investigate bases governing Ca^2+^ binding affinities among three *α*PV variants and (3) understand the factors governing the selectivity of these variants for Ca^2+^ over Mg^2+^. To accomplish these goals, MD simulations and thermodynamic analyses of ion binding were conducted of these *α*PV variants. We discuss factors contributing to ion binding affinity and selectivity in order of importance. Limitations of the Methodology are provided as Supplemental Data (Section S8.5).

### 4.1 Rank-ordering *α*PV variants according to binding affinity

One of our goals was to elucidate the thermodynamic factors governing the variations in affinity among variants of *α*PV. Free energy techniques including thermodynamic integration, umbrella sampling and adaptive biasing force have been used for rigorous estimates of Ca^2+^ affinity for a variety of Ca^2+^-binding proteins [48, 26]. These methods achieve high accuracy, i.e., the estimated free energies are within 1 kcal/mol, but do so with a prohibitive computational expense [49]. Further, it is frequently difficult to decompose free energies predicted by these approaches into per-residue contributions, which limits our ability to implicate specific structural differences between variants to conformations and dynamics to their respective ion affinities. For these reasons, we determined if whether the less accurate, but more computationally-feasible MM/GBSA end-point method [49] could rank-order variants by their relative affinity differences. We found that the default strategy of including Ca^2+^/protein interactions yielded energies inconsistent with the *α*PV proteins’ experimentally measured Ca^2+^ affinities, which we attribute to shortcomings in the evaluation of polar solvation energies (see Limitations). However, our results suggest focusing on the reorganization score of forming the holo state upon ion binding relative to the apo rank-orders the variants correctly.

### 4.2 Contributions of EF-hand loop and co-ordination shell to altered Ca^2+^ binding

The CD and EF domains are the primary binding sites for Ca^2+^ in *α*PV and form the helix-disordered loop-helix EF-hand motif. The disordered loop reorganizes to coordinate its amino acid oxygens with Ca^2+^, which manifests in the reduced RMS fluctuations of all *α*PV variants upon ion binding predicted by our MD simulations. The most significant reductions were apparent within the EF site loop (Fig. 1F-I), which we hypothesize stems from its greater number of glycines (4) compared to the CD site loop (1). This is rationalized by noting that glycine is regarded as the most labile residue of the twenty natural amino acids [50]. Interestingly, we observed that the amplitude of the apo state RMSFs in the Ca^2+^ binding sites correlated with the variants’ ion affinities, with the largest RMSFs reported for S55D/E59D *α*PV and smallest for D94S/G98E *α*PV. The correlation of increased fluctuations in the apo state and Ca^2+^ affinity is consistent with observations of larger Stokes radii and thermal instability for variants of paramecium calmodulin, which were shown to impart increased Ca^2+^ binding affinity relative to the WT [51]. Similarly, Vologzhannikova et al. confirmed the importance of disordered apo states in conferring elevated Ca^2+^ affinity, as they demonstrated *β*PV-1 apo state from coho salmon is unfolded and associated with its increased affinity compared to other *β*PV apo forms that are folded [52]. Especially for the CD-loop sites, the calculated 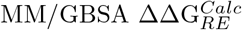 scores clearly indicated that the conformational changes upon Ca^2+^ binding yielded more favorable values for S55D/E59D *α*PV and less favorable for D94S/G98E *α*PV relative to WT(Table S2). Overall, we show that a decreasing apo state stability in intact PVs is associated with increased Ca^2+^ affinity.

It is tempting to interpret differences in affinities among the variants based on properties of just the Ca^2+^ binding loop, as studies have argued that the position and charge of amino acid substitutions can dramatically enhance Ca^2+^ affinity [53]. For instance, it might be expected that increasing the number of oxygens bound to Ca^2+^ could improve affinity, as has been rationalized for selective binding of Ca^2+^ over Mg^2+^ based on bidentate glutamate binding relative to monodentate [21, 22]. Our data instead indicate that the number of protein oxygens predicted to coordinate to the ion is uncorrelated with their affinity. Further, the position of the mutation in the EF-hand loop may also determine the extent of its impact on binding ions. For example, the E101D mutation in PV was found to reduce coordination, which we attribute to the F helix’s limited ability to reorganize to bind Ca^2+^. In contrast, the greater mobility elsewhere in the EF-hand loop facilitated the reorganization of adjacent amino acids to compensate for any loss in coordination due to a given mutation. In addition, we found that bound water can directly participate in coordination through bridging an amino acid with Ca^2+^, which we recently demonstrated was an important factor in modulating Ca^2+^ affinities of the Ca^2+^-binding ATPase SERCA [54]. Despite the lack of correlation between coordination number and affinity, our analyses firmly indicate that loop/ion interactions represent significant contributions to the MM/GBSA-estimated ΔΔ*G*s relative to the WT, which favor S55D/E59D *α*PV over Δ*α*PV. For the most part, the changes in 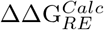 within each loop was dominated by the 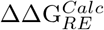 at the mutation site. While the trend in the 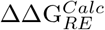 for the sites could be easily rationalized (e.g., polar-to-charged residues like S55D were favorable), the magnitudes of which, especially relative to other mutations, were difficult to surmise without the simulations. This provides evidence that coordination shell alone is just one factor the contributes to Ca^2+^ affinity in *α*PV.

We used these two methods of energetic analysis to quantify the contributions of oxygen and water coordination with Ca^2+^. Using the MM/GBSA analysis approach (Fig. S4) we found that the S55D mutation is thermodynamically unfavorable (11 kcal/mol), whereas the E59D mutation is favorable (−18 kcal/mol), though these calculations neglected stabilizing Ca^2+^/protein interactions. Because of this, we used the MSA approach to estimate ion binding chemical potentials that account for ion binding shell volume, finite sizes of the ions and oxygens, number of coordination oxygens, as well as the ion solvation energy. The MSA approach confirmed that Ca^2+^ was thermodynamically favored over Mg^2+^ across broad binding site volumes and coordination numbers (Fig. 3). Moreover, S55D/E59D *α*PV was shown to more favorably bind Ca^2+^ relative to the WT (Δ*µ* = 0.14 kcal/mol) and D94S/G98E *α*PV (Δ*µ* = 0.21 kcal/mol) on account of its more negative oxygen effective charge (Δ*q*_*eff*_ = *−*0.02*e*). In general, Ca^2+^ affinity had a modest dependence on binding site volumes and coordination numbers. In other words, we speculate that the moderate variations in binding site volumes and oxygen coordination numbers we computed for the *α*PV variants did not appreciably impact the Ca^2+^ chemical potential. In fact, we applied this to a range of PV homologs with at least 50% sequence identity to the WT *α*PV, which included several *β*-parvalbumins structures. The model predicted chemical potentials for a given average effective oxygen charge varied by up to 1 kcal/mol based on the MSA method, despite the diversity of coordination numbers and binding site volumes (see Fig. S8). Instead, the most significant difference was found for PDB 2MBX and 5XND, as the two species have significantly different effective oxygen charges in the binding domain (*−*0.74e versus *−*0.81e). Hence, properties of the ion coordination shell alone appear to have limited ability to rationalize the affinity differences in the *α*PV variants we considered, unless significant changes in the charge density were involved. Even then, consideration of factors like changes in configurational entropy of the loop and strain induced within the loop upon ion binding are likely to significantly influence ion binding thermodynamics [38].

### 4.3 Contributions of protein structure beyond EF-hand loop in determining Ca^2+^ binding

Our analyses of the Ca^2+^/EF-hand binding thermodynamics indicate that there is a strong driving force to optimize Ca^2+^ coordination, which is afforded through the ease with which the EF-hands can reorganize. Although this reorganization appears to normalize Ca^2+^ binding properties within the loops, the respective changes in loop conformations can drive compensatory adjustments to the *α*PV helical bundle. This coupling between EF-hand binding and allosteric changes in bundle organization is exemplified in Ca^2+^ binding proteins such as calmodulin [55, 56], TnC [57, 58] and S100A1 [59, 60, 61]. The openings of these proteins’ hydrophobic peptide-binding domains following Ca^2+^ binding are generally not evident in theMg^2+^-bound complexes. We show that the enhanced binding of Ca^2+^ to *α*PV protein correlates with several structural changes beyond the EF-hand loop region. The predominant changes caused by ion binding include: (1) reduced hydration shell around the protein, (2) increased hydrogen bonding, (3) increased *α* helicity, particularly in the D and E-helix regions, and (4) reduced *β*-strand content between the CD and EF loops. We discuss each of these factors in order of importance based on their effects on the MM/GBSA scores, where applicable.

Our results summarized a variety of changes in binding loop dynamics, helical content, hydrogen bonding, and bundle packing upon ion binding to *α*PV variants that were anticipated to impact the protein/solvent interface. Indeed, we found that the subtle conformational changes induced by ion binding manifested in reduced hydration of holo *α*PV compared to apo and to a similar extent across all variants. This desolvation has been suggested to be a significant factor in driving ion binding in Ca^2+^-selective proteins [22, 21], which can be rationalized by the liberated solvent increasing the configurational entropy of the system. The displacement of loosely-bound waters in the CD and EF loops of the apo states upon binding Ca^2+^ represented a substantial portion of the total numbers of waters liberated in the holo state. It is unclear how much each liberated water contributed to the Ca^2+^ affinity, although one report suggested that a Glu to Asp mutation reduced Ca^2+^ affinity by five fold estimated based on *K*_*D*_*′ /K*_*D*_ = exp(*−*ΔΔ*G/KT*) due to an extra coordinated water [17].

We attributed the remainder of the displaced waters to reorganization of the helical packing that apparently reduced the proteins’ solvent accessible surface area. It has been speculated that the reduced affinity of CBPs for Mg^2+^ relative to Ca^2+^ can be partially attributed to the former ion’s reduced capacity to displace waters [22], but our data showed no significant differences between the ions in terms of protein dehydration. While we were unable to determine the relative thermodynamic contribution of these displaced waters to the MM/GBSA-estimated scores we calculated, the greater number of waters displaced in the S55D/E59D *α*PV and WT cases relative to D94S/G98E *α*PV is consistent with their trends in stronger experimentally-measured binding affinities.

The dampened RMSF values and dehydration of holo-state proteins relative to apo states reported in our simulations implicate changes to the organization of the proteins’ helix bundles. Unlike Ca^2+^ binding proteins such as calmodulin (CaM) that undergo significant conformational changes upon Ca^2+^ binding, the *α*PV proteins exhibit comparatively more subtle changes. To highlight these subtle effects, we measured the net number of hydrogen bonds formed during the apo to holo state transition, as these bonds would be expected to yield larger free energy contributions than corresponding interactions between apolar residues. We found consistent increases in the number of intraprotein hydrogen bonds in the holo states of the S55D/E59D variant and the WT *α*PV compared to Δ*α*PV, which were correlated with the pattern of dehydration discussed in the previous section. Some of this increase was attributed to the extension of the D and E helices’ *α*-helicity, although this increase was generally similar across the variants. Increasing hydrogen bonds generally was observed in the holo versus apo states, with similar changes noted for Ca^2+^ and Mg^2+^. However, longer duration hydrogen bonds appear to be evident in the stronger binders of Ca^2+^, which may contribute to stabilization of the bound complex.

Ca^2+^ binding proteins commonly exert regulatory control through interfacing with target proteins. In many cases, such as for CaM and TnC, the protein-protein interface comprises the CBP’s solvent-exposed hydrophobic patch and a coiled domain from the target peptide. The binding of the target peptide is often associated with an increase in the apparent Ca^2+^ affinity of the CBPs and increased thermostability [18, 62]. The CD and EF helix-loop-helix motifs in the parvalbumin class of CBPs bear high structural similarity to the globular domains of CaM and TnC. Uniquely, the AB domains in PV assume a position similar to that of the target-bound protein-protein interface in CaM and TnC, where it binds its CD and EF domains through seven hydrogen bonds. In this regard, the AB domains in PV serve as an ‘endogenous’ peptide that confers thermostabilty and enhanced Ca^2+^ affinity. Indeed, experimental studies support that this AB domain is essential for stabilizing the apo-state *α*PV into its folded form and protects the *α*PV hydrophobic core from hydration, whereby Ca^2+^ affinity is enhanced [18]. Interestingly, a truncated PV variant that lacks the AB domain maintains the ability to bind Ca^2+^ albeit with significantly reduced affinity, whereas a similar truncation in the structurally-similar calbindin 9K is incapable of binding the ion [62].

Our simulation results confirm that the intact *α*PV has increased hydrogen bonding and reduced bundle mobility as evidenced by RMSF calculations, helicity, and dehydration upon Ca^2+^ binding relative to the truncated variant. The differences highlight the importance of the AB domains in defining *α*PV’s Ca^2+^ binding properties. Supporting these observations, in the holo state there is a loss of favorable reorganization score of the CDEF domain in the truncated variant compared to the intact one (Table 1). Despite this loss of stability, the Δ*α*PV holo state remains folded in the presence of Ca^2+^ as supported by the Thepaut *et al* experiments [18]. Nonetheless, even though the Δ*α*PV holo structure is folded, it presents high RMSF values that suggest instability within the helix bundle relative to the intact *α*PVvariants, in addition to reduced D-helicity and enhanced *β*-sheet character that may impair Ca^2+^ binding. While we were unable to predict the unfolded apo state indicated by Thepaut *et al*, our simulations suggest that the Δ*α*PV apo structures were unfolding and highly labile relative to the intact *α*PV structure. From this perspective, the presence of the AB domains may amortize the thermodynamic cost of folding the protein upon binding Ca^2+^, which would otherwise reduce the *α*PV apparent Ca^2+^ affinity. In support of this interpretation, lysine-to-asparagine mutations of the AB domain sites A8 and H26 in PV constructs modeled after Antartic fish were found to increase Ca^2+^ binding affinity and was attributed to forming additional hydrogen bonds within the AB or between AB and CDEF domains [63]. Overall, our observations highlight the importance of AB/CDEF domain interactions for maintaining high Ca^2+^ binding affinity.

As with many Ca^2+^ binding proteins, *α*PV is nearly entirely composed of *α*-helices that undergo displacements and changes in folding upon binding by Ca^2+^ [64, 65, 66, 22]. We observed a consistent trend in our simulations of increased *α*-helicity in the ion-bound holo states relative to apo, with greater gains demonstrated for Ca^2+^ relative to Mg^2+^. This observation is consistent with observations from Laberge et al, who through ultraviolet circular dichroism spectra and picosecond MD simulations of cod PV [67] showed that removal of Ca^2+^ reduced *α*-helical content. Our simulations indicate that the majority of the gain in *α*-helical character stems from refolding of the N-terminal end of the D helix. At these termini, glutamic acids at the 59th and 62nd sequence positions (E59D for S55D/E59D *α*PV) are reoriented by Ca^2+^ as the ion binds the carboxylic acid side chains. This process increases the helicity of residues E/D59 through G64 to 90%. In contrast, we observed increase in helicity in the C-terminal end of the E helix, particularly for the residues T84-D90.

Interestingly, several observations are notable: firstly, Ca^2+^-driven changes in D-helix *α* character were limited to the N-terminus up to G64. Glycine is frequently found to disrupt or break *α*-helices [68, 50], thus we speculate that varying the relative position of this residue within the D-helix could modulate Ca^2+^ affinity (see Section S8.4 for further discussion); second, our Δ*α*PV simulations reflected a very low (55%) degree of helicity formed in the D-helix N-terminus, which suggests the AB domains likely play an important role in stabilizing the Ca^2+^ binding loop, which in turn may increase its affinity for Ca^2+^; similarly, weaker binding *α*PV variants tend to have less helical character in their apo states. Lastly, the D-helix N-terminus has a lesser degree of *α*-helicity in the Mg^2+^-bound state (65%). It has been speculated that this difference may be the consequence of the 12th position glutamic acid (E62) promoting bidentate oxygen coordination for Ca^2+^ relative to monodentate for Mg^2+^ [22, 37]. No such differences were noted in the E-helix between the Ca^2+^ and Mg^2+^ ions. Altogether, our data indicate that increasing the *α*-helicity particularly in the N-terminal region of helices D and E, may play an integral role in high affinity Ca^2+^ binding and selectivity against Mg^2+^.

A hallmark of Ca^2+^ binding proteins is the pairing of two Ca^2+^-binding EF hand domains via a *β*-sheet, which has been coined as the “EF-hand *β*-scaffold (EFBS)” [22]. These *β*-sheets are believed to contribute to the Ca^2+^ binding mechanism and cooperativity for some CBPs [22]. Our studies support the notion that *β*-sheet character is reduced upon Ca^2+^ binding and to a lesser extent by Mg^2+^. Among the intact *α*PV variants, the reduction correlates with Ca^2+^ affinity and analogously, the low Ca^2+^-affinity Δ*α*PV variant presents the highest degree of *β*-sheet character when bound to Ca^2+^. It is evident from published structural data and simulations that the reduction in *β*-strand content occurs as the residues contributed by each site (F57-E59 and K96-G98) are drawn away from the *β*-sheet interface by their respective coordinated ions. The largest displacements arise from E59 and K96 as they are separated from one another by S55 and D94, respectively, through the formation of hydrogen bonds (Fig. S9). Our simulation data indicate that the coordination geometries and extent of *β*-sheet character were similar among the holo variants, whereas the apo state *β*-sheet content increased for variants with reduced Ca^2+^ affinity. Hence, the reduced binding affinity for Ca^2+^ in those variants likely reflects the thermodynamic cost of disrupting a greater number of hydrogen bonds participating in the *β*-sheet interface. Interestingly, it has been reported that the coupling between the two binding sites through hydrogen bonds and dipole-dipole interactions improved Ca^2+^ affinity to the CBPs through positive cooperativity [20], although the lack of cooperativity in *α*PV may give rise to this contradictory trend in our data. Overall, our simulation data suggest that mutations that enhance *β*-sheet character in the apo state appear to offset the free energy gain upon Ca^2+^ binding.

### 4.4 Molecular determinants of Ca^2+^ versus Mg^2+^

*α*PV selects for Ca^2+^ over the smaller Mg^2+^ ion that is roughly 10000-fold more concentrated within cells [22]. The experimentally-determined affinity difference of *≈* 10 kcal/mol in large part can be rationalized by the considerably higher desolvation cost for Mg^2+^ relative to Ca^2+^ and to a lesser extent by the reduced number of protein oxygens bound by Mg^2+^ [22], as was confirmed by our MSA calculations. Additional factors, however, though can further tune the observed affinity differences for these ions, including the energetic cost of distorting the EF-hand to accommodate the smaller Mg^2+^ ion over Ca^2+^ [37], polarizability [69, 70] and many-body polarization effects [26], which depend on the number and the arrangement of charged residues. Regardless of the origins of the affinity differences, our simulations reveal distinct differences between the holo states of Mg^2+^-bound *α*PV structures relative to those bound with Ca^2+^. These differences include reduced *α*-helicity, especially within the D-helix N-terminal region, and less significant disruption of the *β*-sheets bridging the EF-hand loops. These secondary structure differences are mirrored by the MM/GBSA scores from these corresponding regions in the *α*PV proteins. Similarly, it could be argued that those differences may have significant impact on proteins such as TnC, for which modest differences in EF-hand structures when bound to Mg^2+^ versus Ca^2+^ dictate the exposure of hydrophobic surfaces necessary for binding peptide substrates. In contrast, the presence of the AB domains in *α*PV most likely masks such structural differences that would arise for Ca^2+^-bound states relative to Mg^2+^ bound. Instead, the AB domains appears to lock *α*PV into an open-like state resembling that of TnC even in the absence of Ca^2+^, which thereby facilitates binding of other competing cations such as Mg^2+^. This interpretation is supported by studies from Thepaut *et al* that indicated Mg^2+^ was unable to fold the Δ*α*PV variant that lacked the AB helices. For this reason, the prominence of parvalbumin proteins serving as delayed Ca^2+^ buffers in skeletal and cardiac muscle [71] likely stems from its AB domain facilitating Mg^2+^ binding and thereby competing with Ca^2+^ binding.

## 5 Conclusions

In this report, we investigated several structural features in *α*PV that factor into the proteins’ affinity for Ca^2+^ and its selectivity over Mg^2+^. Our approach utilized molecular dynamics simulations of the wild-type *α*PV ion-free (apo) and Ca^2+^/Mg^2+^-bound (holo) states, in addition to simulations of *α*PV variants with increased or reduced Ca^2+^ affinity relative to the WT. We demonstrate that the MM/GBSA endpoint method (when applied to the protein reorganization score) and MSA theory are effective tools for qualitatively rank ordering the intact *α*PV variants (WT, S55D/E59D *α*PVand D94S/G98E *α*PV) by their experimentally determined affinities. We used scores from these analyses to then implicate structural changes that rationalize the variants’ ion binding affinities. We find that simple metrics such as the number of coordinating oxygens about a bound ion or changes in the binding sites’ net charge are insufficient to completely rationalize observed changes in affinity. Rather, concurrent effects on the protein’s secondary structure composition, solvation properties, and hydrogen bonding networks throughout its globular domain are involved and must be carefully analyzed to understand the effects of mutations and evolutionary differences between PV isoforms. Clearly, rationalizing affinity in Ca^2+^ binding proteins including PV therefore necessitates not only knowledge of the proteins’ primary sequence and corresponding structures, but also how those structures impact the thermodynamics of the Ca^2+^ free and bound states. In many cases, this may require looking beyond direct effects of changing the charge or Ca^2+^ coordination within the EF hands [72]. This firmly indicates the value of molecular simulations in probing residue-specific contributions to protein stability and Ca^2+^ binding in *α*PV. Thus, it is apparent from our data that point mutations can modulate affinity using diverse mechanisms.

An important outcome of our study are potential mechanisms that can be manipulated through mutagenesis to tune Ca^2+^ affinity. As an example, introducing mutants that destabilize the *β*-strands linking the CD and EF hand loops in the apo state may amortize their reorganization when binding Ca^2+^. Additional strategies could include increasing the apo state *α*-helicity by replacing the helix-breaking G64 with a small hydrophobic residue like alanine (G64A) that would restore normal helicity, introducing additional hydrogen-bonding polar residues to bridge the AB and CDEF interface and thereby tighten AB/CDEF binding, as well enriching the CD and EF loops with glycine to further increase their mobility.

## 6 Materials and methods

### 6.1 Protein systems

Ca^2+^ bound rat *α*PV X-ray crystal structures (PDBs: 1RWY [40], 1XVJ [16], 1S3P [15], and 1G33 [18]) were used for the MD simulations. Apo simulations were set up by removing ions from the holo crystal structures. Because crystal structures of Mg^2+^ bound and apo *α*PV protein are not yet available, Ca^2+^ was replaced with Mg^2+^ for the Mg^2+^ bound simulations. System preparation was performed with *tleap* from Amber 16 [73].

### 6.2 Classical MD

All systems were solvated in TIP3P [74] water box with 20 Å margin and neutralized with 0.15 M KCl. Each system has about 14,000 waters, 82 neutralizing ions, for a total of *≈*44,000 atoms. The systems were parameterized with the amber ff12SB force field [75] with Ca^2+^ and Mg^2+^ parameters were adapted from Li-Merz [29]. The system was subjected to 6000 steps minimization with protein atoms being fixed during the first 1000 steps. The system was then heated under NVT ensemble to 300 K over 0.1 ns using the weak-coupling algorithm [76]. Constraints were introduced on protein atoms with force constant as 500 kcal/mol.Å^2^. The heated system was equilibrated for 1 ns under the NPT ensemble without constraints on the protein. After equilibrium, each system was simulated in triplicate for *≈*1 *µ*s under the NPT ensemble at 300 K temperature. A 2 fs time step was used with the non-bonded cutoff set as 10 Å and the electrostatic interactions were treated the with Particle Mesh Ewald (PME) method [77]. The SHAKE algorithm was used [78]. The simulation data of the production run were saved for every 20 ps. The minimization and equilibration were performed using the *PMEMD* module while the production runs were performed using the GPU-accelerated *PEMD*.*CUDA* module of AMBER16 [73].

### 6.3 Analyses

Two snapshots for each ns of trajectory were considered for the analysis (*≈*2000 data points for each simulation trajectory (*≈*1*µ*s) and a total of 72000 data points for all systems combined (12 systems *×* 3 trials *×* 2000 data points/ 1*µ*s) *≈*72000 data points). Almost all analyses were conducted with VMD [79] and its associated plugins [79, 80], except for the radial distribution functions. For ion bound systems, the radial distribution of chelating oxygens about the cation (either Ca^2+^ or Mg^2+^) was obtained by the *radial* command of *CPPTRAJ* [81]. For calculating hydrogen bonds, the cut-off distance and angle used were 3.5Å and 30*°* respectively. RMSDs and RMSFs were calculated by superposing each trajectory frame with the respective x-ray crystal structure. The MMPBSA.py script of AMBER16 was used for the MM/GBSA calculations. The ionic strength for the MM/GBSA calculation was set as 0.15 M with the generalized Born model option set as igb=5. The entropy was neglected during the MM/GBSA calculations. The entire MD trajectories were used for the MM/GBSA calculations. The total score/reorganization score of each system was calculated as the sum of all five individual components of the MM/GBSA method including electrostatic, van der Waals, internal, polar and non-polar solvation, respectively.

### 6.4 MSA analyses

The mean field approximation theory (MSA) self-consistently solves for the optimal distribution of ions comprising an electrolyte solution within a protein binding domain, or ‘filter’, consisting of mobile oxygens. To do this, the model utilizes a Hamiltonian reflecting the free energy of packing finite-sized ions as hard spheres and electrolyte-shielded electrostatic energies in a fixed volume of oxygens that cannot exchange with the surrounding medium [46, 82]. This method includes the thermodynamics of solvent ions simultaneously competing with the binding site, the desolvation energies of those ions, as well as the finite sizes of ions and oxygens in the binding domain. Ion distributions, *ρ*, are predicted from the chemical potential, *µ*, which arises from the balance between (un)favorable electrostatic interactions and entropic hard-sphere interactions [83].

All ions, namely Ca^2+^, Mg^2+^, K^+^ and Cl^*−*^ can readily repartition in order to minimize the gradient in the chemical potential:

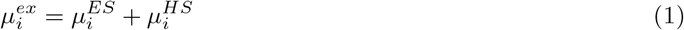

Further details of the forms of 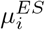 and 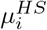 are provided in the supplementary material of [38]. At steady state, for each ion species, the following equilibration is reached:

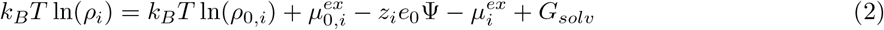

where *ρ*_*i*_ is the density of the i-th ion type in the binding site, *ρ*_0,*i*_ is the density of the i-th ion in the bath (assumed to remain constant), 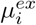 is the excess chemical potential of the i-th ion in the binding site, 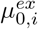 is the excess chemical potential of the i-th ion in the bath (also constant), *z*_*i*_ is the integer charge of the i-th ion species, Ψ is the Donnan potential for the binding site, *G*_*solv*_ is the solvation energy of the i-th ion.

## 7 Acknowledgements

Research reported in this publication, release was supported by the Maximizing Investigators’ Research Award (MIRA) (R35) from the National Institute of General Medical Sciences (NIGMS) of the National Institutes of Health (NIH) under grant number R35GM124977. PKH thanks Caitlin Scott, Charles Adeniran and Andrew Mondragon for prototyping simulation conditions for this project. We also thank XSEDE for providing resources for conducting the simulations [84].

## S8 Supplement

### S8.1 Figures

**Figure S1:**
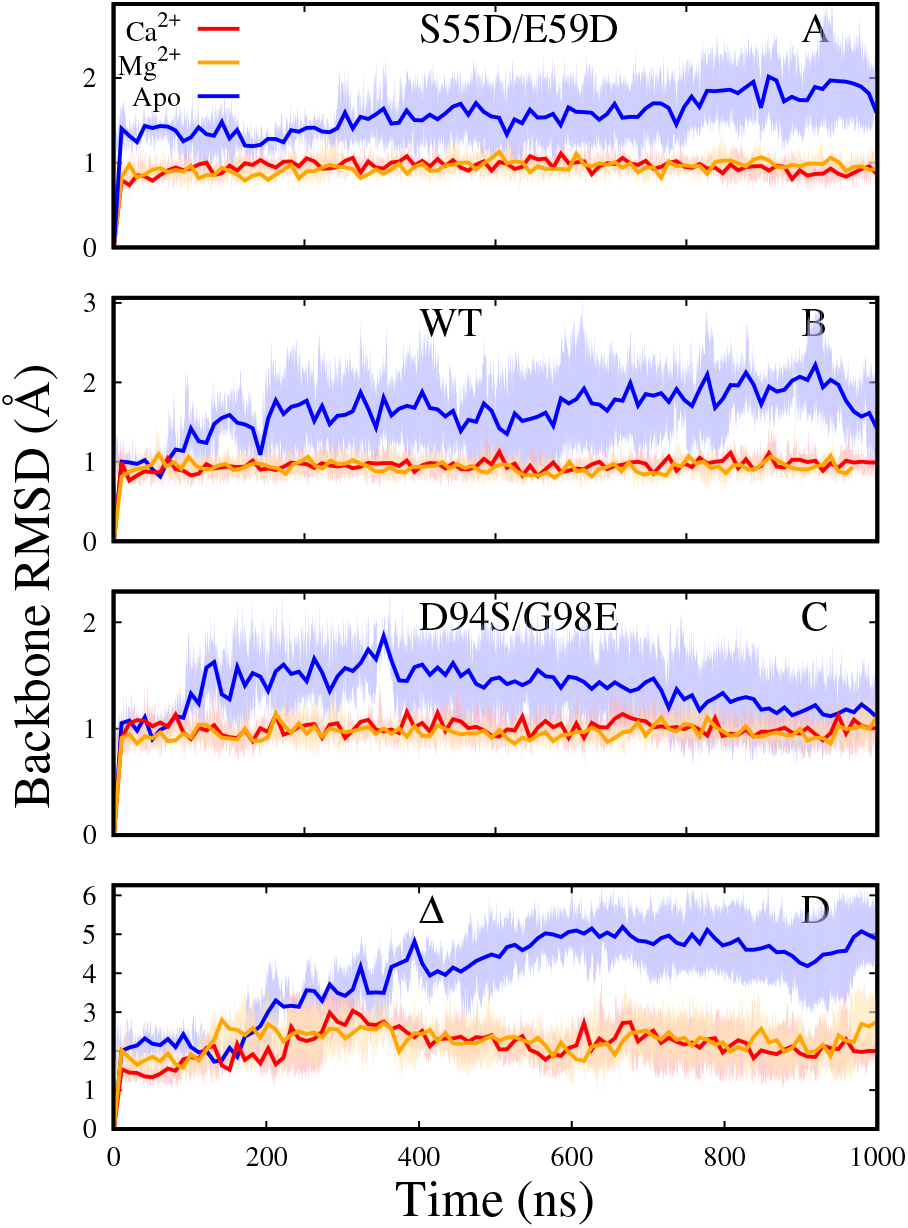
Moving average of the backbone RMSD for S55D/E59D *α*PV, *α*PV, D94S/G98E *α*PV and Δ*α*PV protein. Apo, Ca^2+^, and Mg^2+^bound systems are colored blue, red and orange respectively. Dark lines represent the average and shaded regions represent the standard deviation of three MD runs.

**Figure S2:**
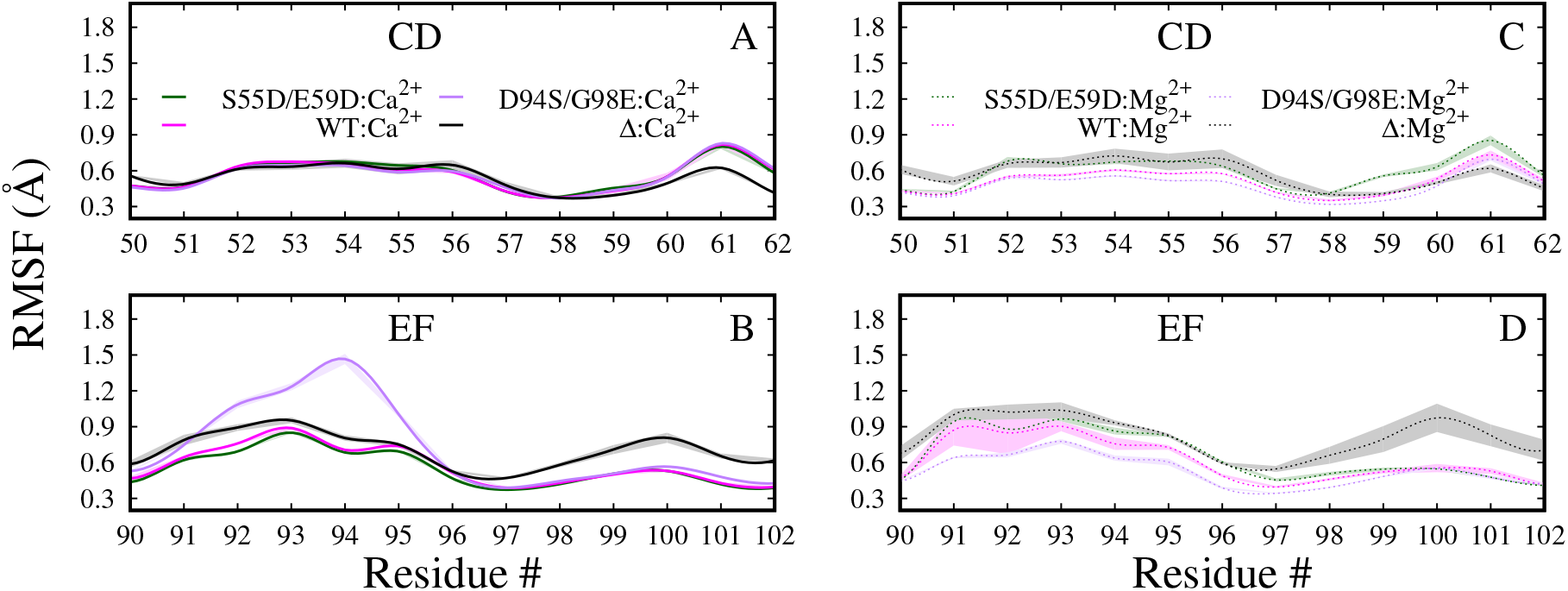
Root mean square fluctuation (RMSF) of the CD (A, C) and EF (B, D) sites of holo systems. Ca^2+^ bound sytems are shown in panels A and B and Mg^2+^ bound in the panels C and D. Mean and standard deviation of three MD trials are shown.

**Figure S3:**
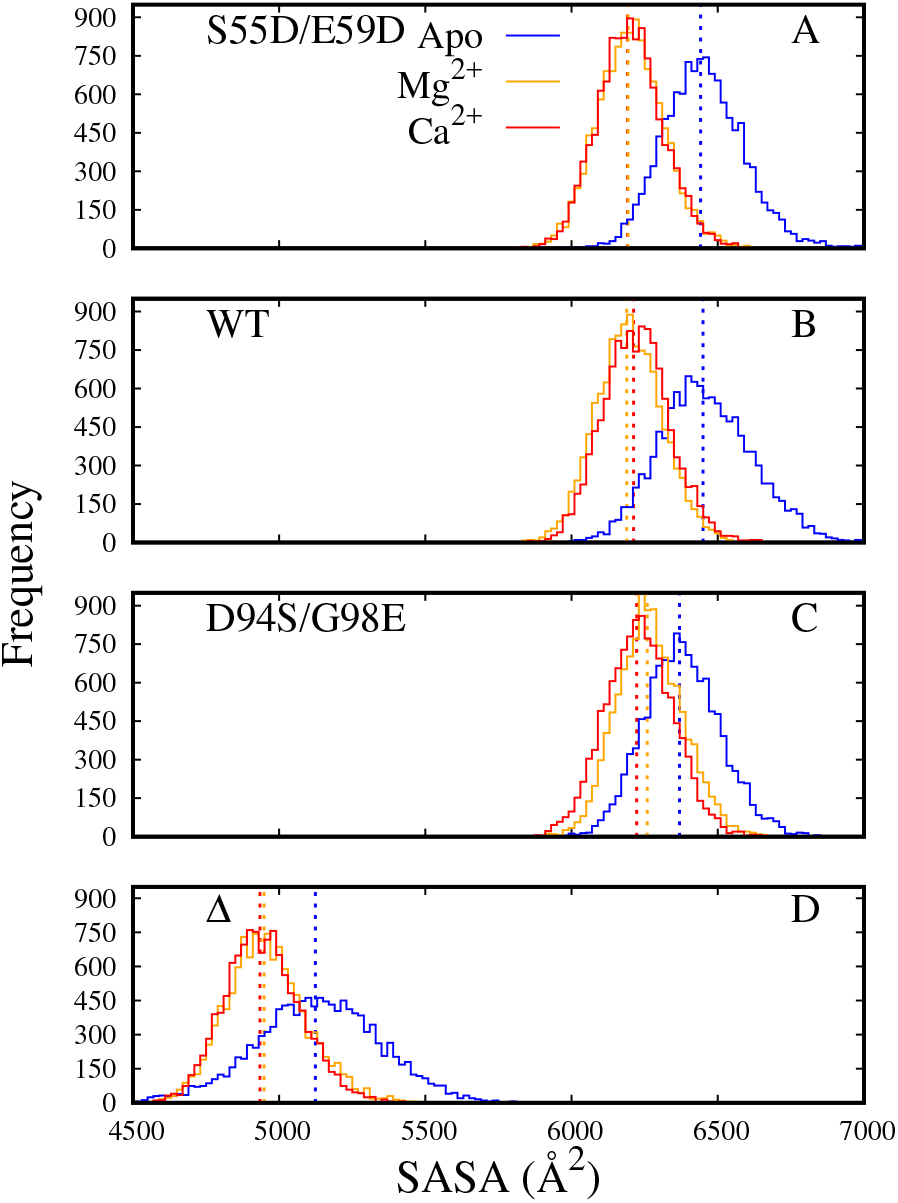
Distributions of solvent accessible surface area (SASA) of entire protein. Apo, Ca^2+^, and Mg^2+^bound systems are colored blue, red and orange respectively. Averages of all three trials for each *α*PV variant are represented by dashed vertical lines..

**Figure S4:**
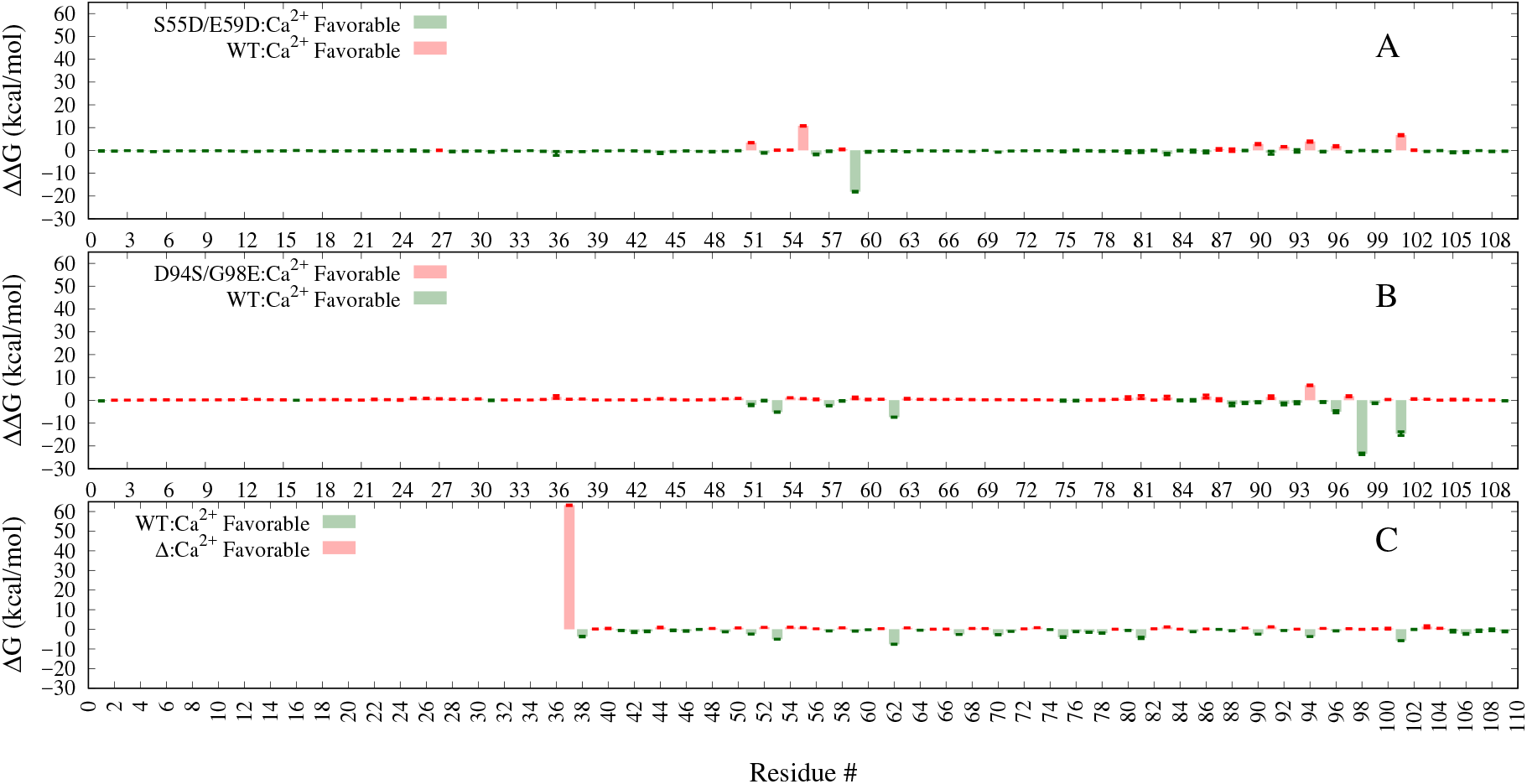
Residue reorganization energies of all Ca^2+^ bound systems calculated using MM/GBSA. The MD predicted Δ*α*PV apo does not unfold as expected from experiment [18], hence only free energies of holo state are compared. See Table S3 and Table S4 for summations of structurally-significant regions. Mean and sem of each residue estimated from three MD trials are shown.

**Figure S5:**
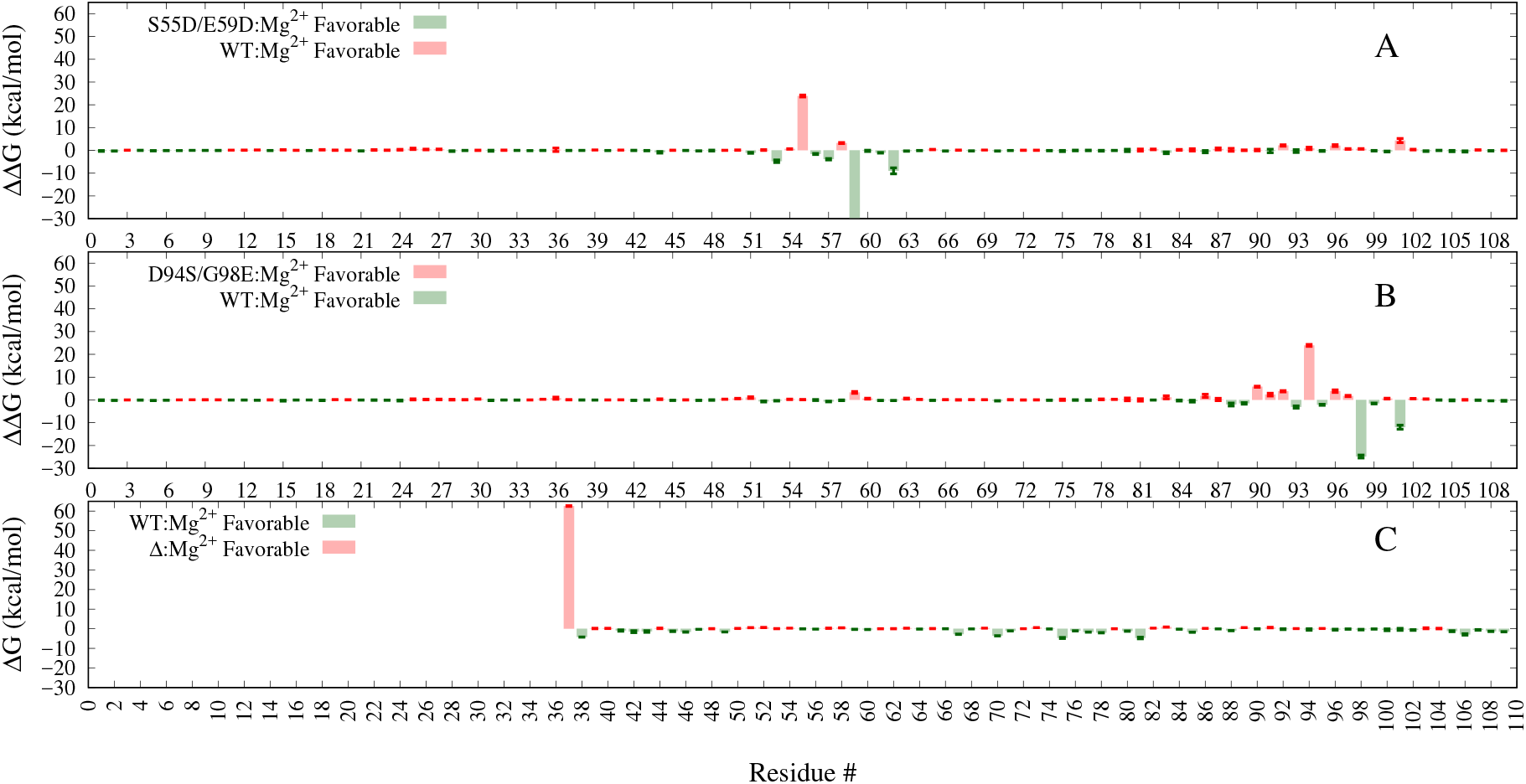
Comparison of MM/GBSA residue reorganization energies of all Mg^2+^ bound systems. Mean and sem of each residue estimated from three MD trials are shown.

**Figure S6:**
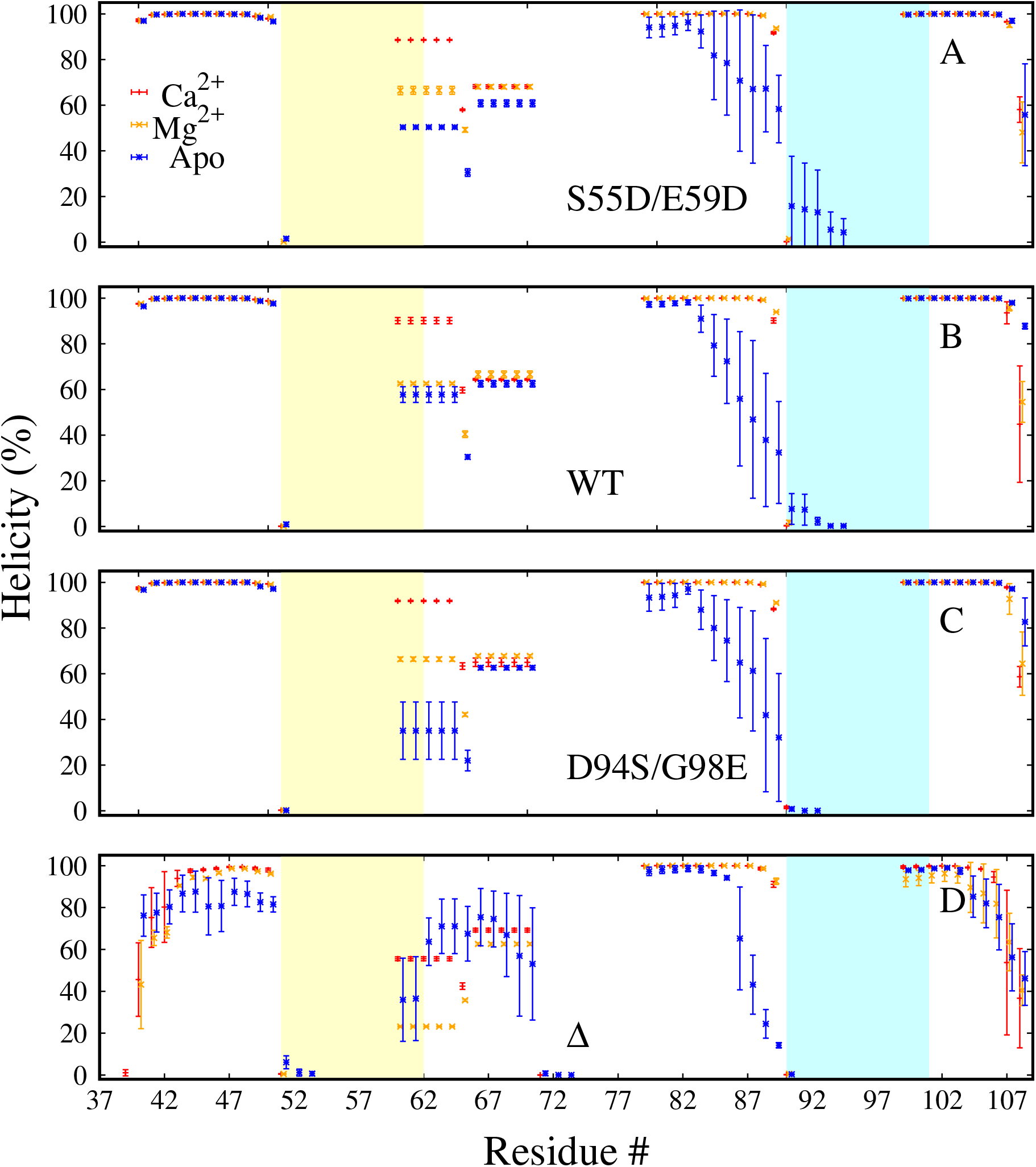
*α*-helical content versus residue number. Apo, Ca^2+^ and Mg^2+^ bound systems are colored blue, red and orange respectively. CD and EF site loop domains are colored yellow and blue respectively. Error bars represent standard deviation estimated from three MD runs.

**Figure S7:**
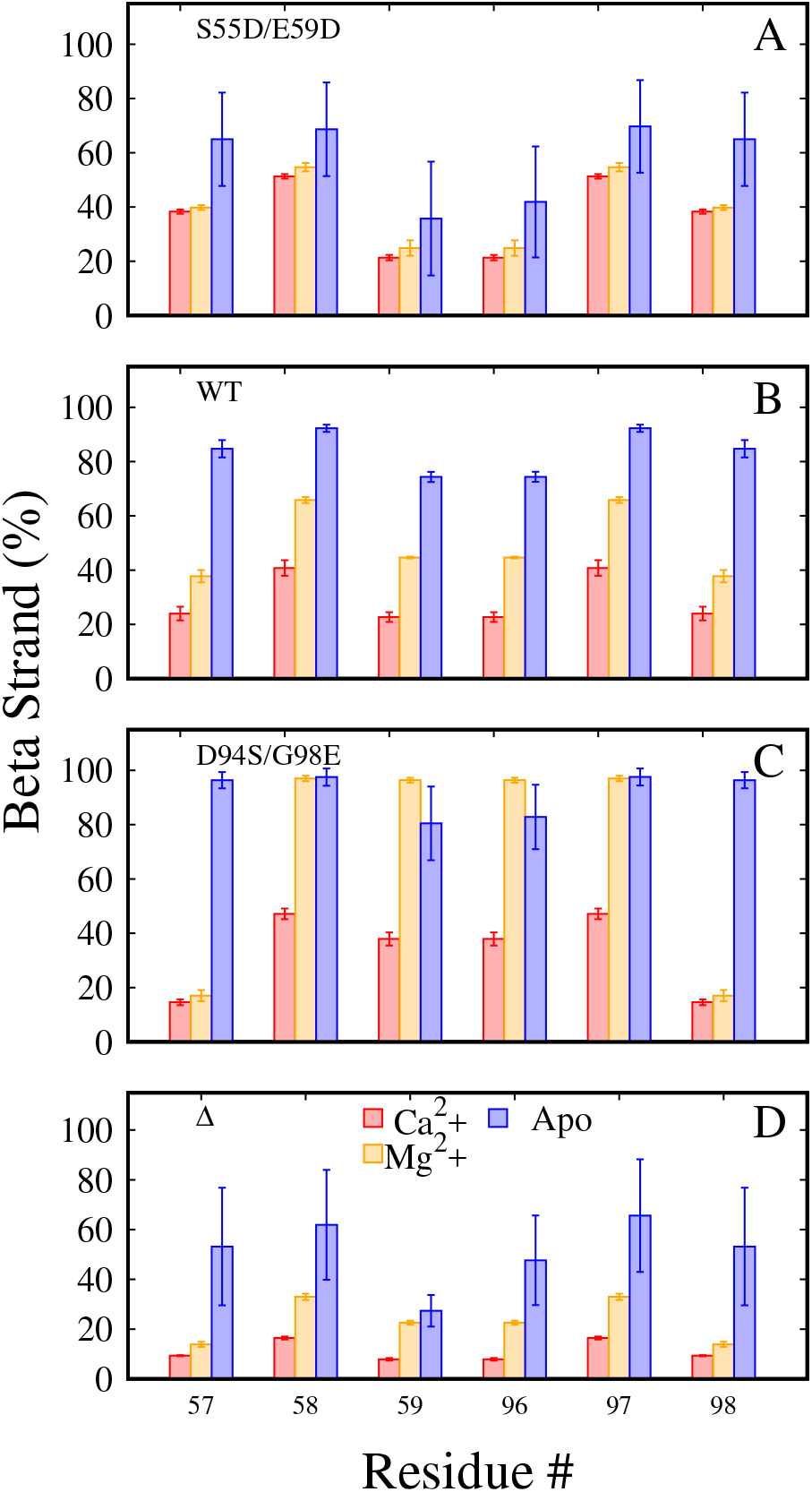
*β*-strand (A-D) of the six residues (F57-E59 and K96-G98) that form the *β*-*β* bridge coupling the two ion binding sites. Apo, Ca^2+^ and Mg^2+^ bound systems are colored blue, red and orange respectively. Error bars represent standard deviation.

**Figure S8:**
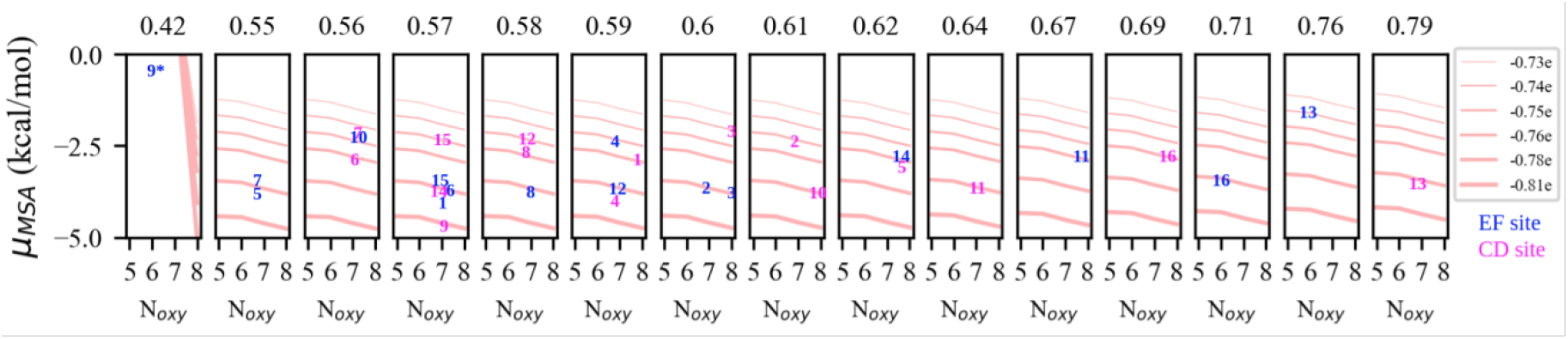
Mean spherical approximation (MSA) calculated Ca^2+^ potentials using binding site information from PDB structures of PV isoforms that have above 50% identity with *α*PV (PDB 1RWY). The identities were calculated via the protein BLAST tool from the NCBI website using the amino acid sequence of PDB 1RWY as input. These PV isoforms are: 1-whiting PV (PDB 1A75), 2-*β*PV (PDB 1B8R), 3-*β*PV (PDB 1B9A), 4-silver hake PV (PDB 1BU3), 5-carp PV (PDB 1CDP), 6-oncomodulin (PDB 10MD), 7-pike PV (PDB 1PAL), 8-pike PV (PDB 1PVA), 9-gadus PV (PDB 2MBX), 10-rat *α*PV (PDB 3F45), 11-gallus *β*PV (PDB 3FS7), 12-leopard shark PV (PDB 5PAL), 13-pacific mackerel PV (PDB 5XND), 14-mustelus griseus PV (PDB 5ZGM), 15-mustelus griseus PV (PDB 5ZGM) and 16-*α*PV(PDB 1RK9). *The data for 2MBX EF site is not included as the cation-oxygen distance is suspiciously small.

**Figure S9:**
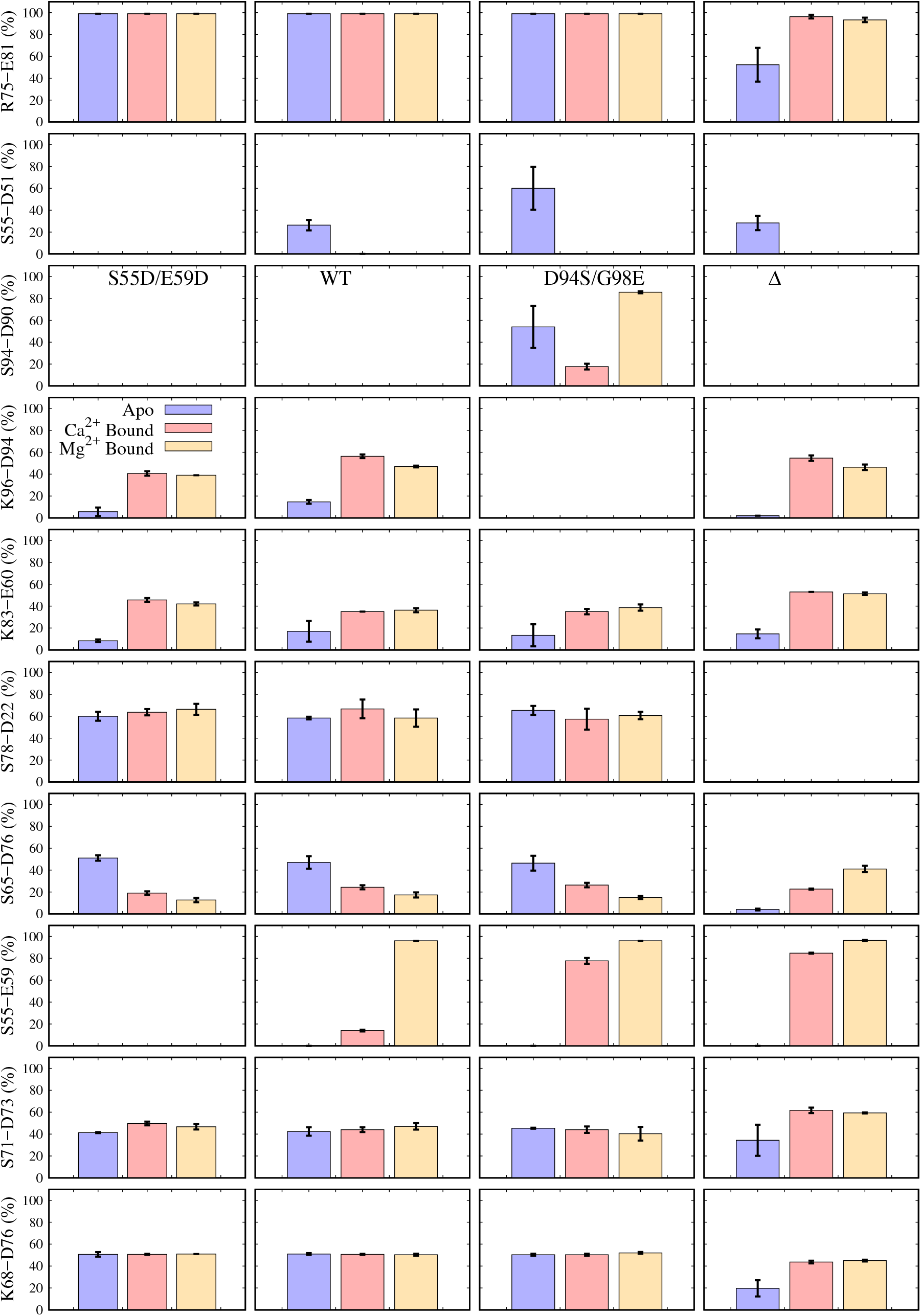
Comparison of specific hydrogen bonds (sidechain-sidechain). Columns 1, 2, 3 and 4 represent *α*PV, S55D/E59D *α*PV, D94S/G98E *α*PV and Δ*α*PV systems respectively. Blue, red and orange represents apo, Ca^2+^ and Mg^2+^ bound systems respectively. Error bars represent standard deviation.

### S8.2 Tables

**Table S1:**
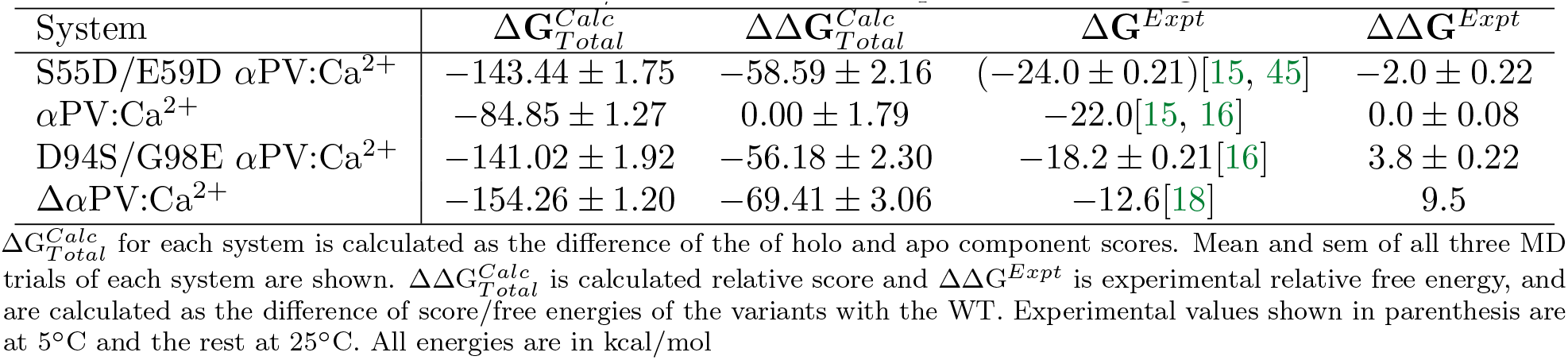
Estimated MM/GBSA scores and experimental binding affinities.

**Table S2:** MM/GBSA reorganization energies of CD and EF site residues.

| System | $G_{RE}^{Calc}$ (CD) | $\Delta G_{RE}^{Calc}$ (CD) | $\Delta\Delta G_{RE}^{Calc}$ (CD) | $G_{RE}^{Calc}$ (EF) | $\Delta G_{RE}^{Calc}$ (EF) | $\Delta\Delta G_{RE}^{Calc}$ (EF) | $\Delta\Delta G_{RE}^{Calc}$ (Total) |
| --- | --- | --- | --- | --- | --- | --- | --- |
| S55D/E59D αPV:Ca<sup>2+</sup> | | $59.13 \pm 1.23$ | $-7.83 \pm 1.33$ | | $57.66 \pm 0.17$ | $13.96 \pm 0.90$ | $6.13 \pm 1.61$ |
| αPV:Ca<sup>2+</sup> | $(-265.86 \pm 0.06)$ | $66.96 \pm 0.52$ | $0.00 \pm 0.74$ | $(-248.05 \pm 0.09)$ | $43.70 \pm 0.88$ | $0.00 \pm 1.25$ | $0.00 \pm 1.46$ |
| D94S/G98E αPV:Ca<sup>2+</sup> | | $79.60 \pm 1.01$ | $12.64 \pm 1.14$ | | $81.94 \pm 1.87$ | $38.24 \pm 2.07$ | $50.8 \pm 2.37$ |
| $\Delta\alpha$PV:Ca<sup>2+</sup> | $(-254.13 \pm 0.04)$ | $79.18 \pm 1.32$ | $12.22 \pm 1.42$ | $(-237.57 \pm 0.37)$ | $55.90 \pm 1.08$ | $12.20 \pm 1.39$ | $24.42 \pm 1.99$ |
| | | $(11.73 \pm 0.07)$ | | | $(10.48 \pm 0.38)$ | | |
| S55D/E59D αPV:Mg<sup>2+</sup> | | $117.01 \pm 2.70$ | $-25.26 \pm 2.76$ | | $116.75 \pm 1.13$ | $9.59 \pm 1.65$ | $-15.67 \pm 3.22$ |
| αPV:Mg<sup>2+</sup> | $(-190.54 \pm 0.21)$ | $142.28 \pm 0.56$ | $0.00 \pm 0.80$ | $(-184.60 \pm 0.82)$ | $107.15 \pm 1.20$ | $0.00 \pm 1.70$ | $0.00 \pm 1.88$ |
| D94S/G98E αPV:Mg<sup>2+</sup> | | $138.79 \pm 1.01$ | $-3.49 \pm 1.16$ | | $108.21 \pm 1.67$ | $1.05 \pm 2.06$ | $-2.43 \pm 2.37$ |
| $\Delta\alpha$PV:Mg<sup>2+</sup> | $(-192.21 \pm 0.04)$ | $141.10 \pm 1.32$ | $-1.17 \pm 1.44$ | $(-182.21 \pm 1.16)$ | $111.26 \pm 1.53$ | $4.10 \pm 1.95$ | $2.92 \pm 2.43$ |
| | | $(-1.67 \pm 0.21)$ | | | $(2.31 \pm 1.42)$ | | |

**Table S3:** MM/GBSA reorganization energies of *β*-sheet residues.

| System | $G_{RE}^{Calc}(\beta)$ | $\Delta G_{RE}^{Calc}(\beta)$ | $\Delta\Delta G_{RE}^{Calc}(\beta)$ |
| --- | --- | --- | --- |
| S55D/E59D αPV:Ca<sup>2+</sup> | | $-4.32 \pm 0.55$ | $-16.78 \pm 0.68$ |
| αPV:Ca<sup>2+</sup> | $(-30.24 \pm 0.15)$ | $12.45 \pm 0.39$ | $0.00 \pm 0.56$ |
| D94S/G98E αPV:Ca<sup>2+</sup> | | $40.84 \pm 0.37$ | $28.38 \pm 0.55$ |
| $\Delta\alpha$PV:Ca<sup>2+</sup> | $(-28.84 \pm 0.06)$ | $14.89 \pm 0.64$ | $2.43 \pm 0.75$ |
| | | $(1.40 \pm 0.16)$ | |
| S55D/E59D αPV:Mg<sup>2+</sup> | | $14.60 \pm 0.59$ | $-28.74 \pm 0.72$ |
| αPV:Mg<sup>2+</sup> | $(0.64 \pm 0.17)$ | $43.34 \pm 0.40$ | $0.00 \pm 0.57$ |
| D94S/G98E αPV:Mg<sup>2+</sup> | | $60.19 \pm 0.55$ | $16.85 \pm 0.68$ |
| $\Delta\alpha$PV:Mg<sup>2+</sup> | $(1.22 \pm 0.17)$ | $44.96 \pm 0.66$ | $1.62 \pm 0.78$ |
| | | $(-1.06 \pm 0.24)$ | |

Data are included for the $\beta$ -sheet residues (F57-E59, K96-G98). Error bars represent sem. Holo state energies are shown in parenthesis.

**Table S4:** MM/GBSA reorganization energies of the D-helix residues.

| System | $G_{RE}^{Calc}(D)$ | $\Delta G_{RE}^{Calc}(D)$ | $\Delta\Delta G_{RE}^{Calc}(D)$ |
| --- | --- | --- | --- |
| S55D/E59D αPV:Ca<sup>2+</sup> | | $23.79 \pm 0.29$ | $-3.76 \pm 0.35$ |
| αPV:Ca<sup>2+</sup> | $(-149.00 \pm 0.17)$ | $27.55 \pm 0.18$ | $0.00 \pm 0.26$ |
| D94S/G98E αPV:Ca<sup>2+</sup> | | $31.18 \pm 0.41$ | $3.62 \pm 0.45$ |
| $\Delta\alpha$PV:Ca<sup>2+</sup> | $(-137.67 \pm 0.14)$ | $34.45 \pm 1.83$ | $6.89 \pm 1.84$ |
| | | $(11.33 \pm 0.22)$ | |
| S55D/E59D αPV:Mg<sup>2+</sup> | | $32.23 \pm 1.27$ | $-11.1 \pm 1.27$ |
| αPV:Mg<sup>2+</sup> | $(-133.21 \pm 0.08)$ | $43.35 \pm 0.11$ | $0.00 \pm 0.16$ |
| D94S/G98E αPV:Mg<sup>2+</sup> | | $42.65 \pm 0.36$ | $-0.71 \pm 0.38$ |
| $\Delta\alpha$PV:Mg<sup>2+</sup> | $(-126.82 \pm 0.06)$ | $45.29 \pm 1.82$ | $1.94 \pm 1.82$ |
| | | $(6.39 \pm 0.10)$ | |

Residues E60 to F70 are considered for this calculation. All energies in kcal/mol. Holo state energies are shown in parenthesis. Error bars represent sem.

**Table S5:**
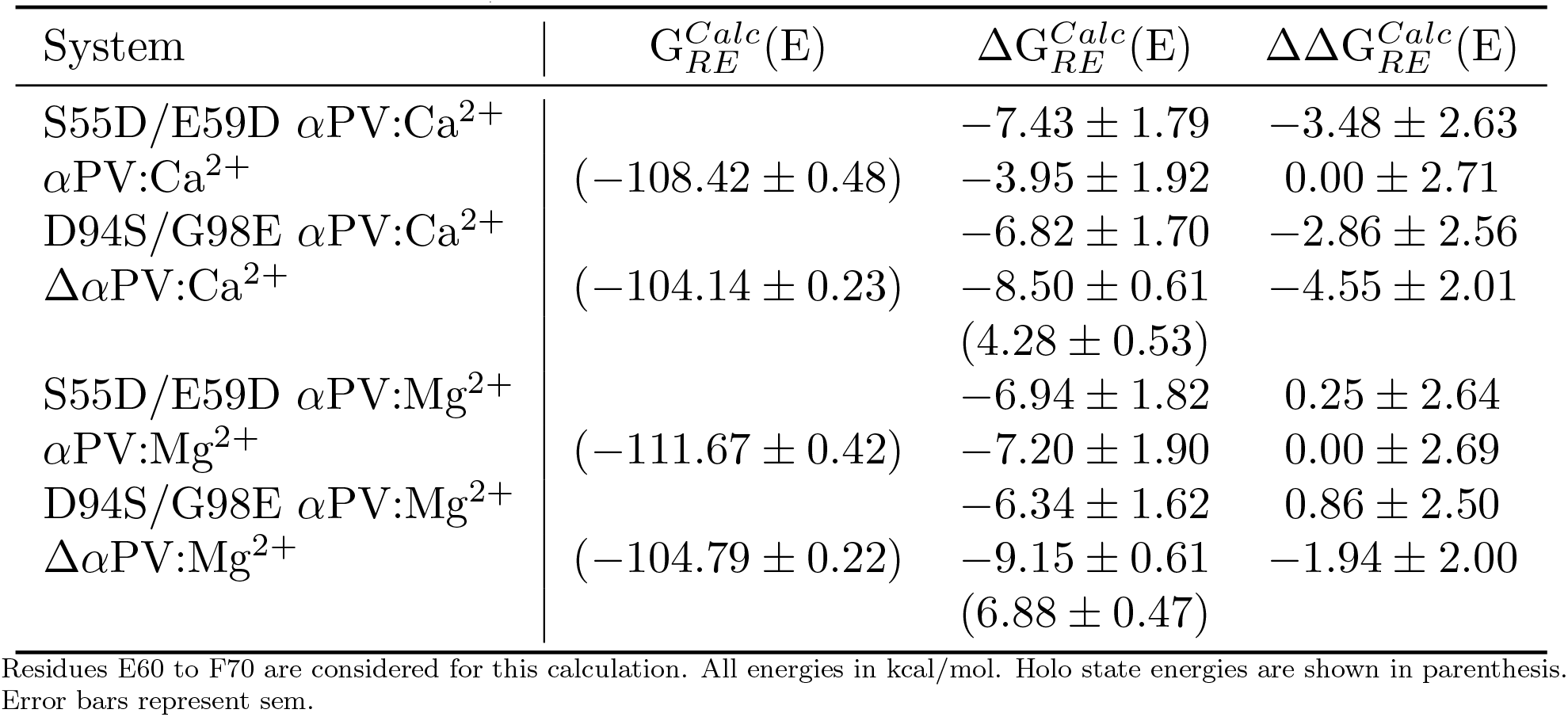
MM/GBSA reorganization energies of the E-helix residues.

**Table S6:** Estimate of effective oxygen charge (q_*eff*_) for each binding site configuration based on MD simulation.

| Case | Loop | Backbone (-0.56e) | Hydroxyl (-0.65e) | Carbonyl (-0.81e) | Water (-0.83e) | $q_{eff}$ |
| --- | --- | --- | --- | --- | --- | --- |
| S55D/E59D | CD | 1.0 | 0.0 | 5.0 | 1.0 | -0.77e |
|  | EF | 1.0 | 0.0 | 5.0 | 1.0 | -0.77e |
| αPV | CD | 1.0 | 1.0 | 5.5 | 0.0 | -0.75e |
|  | EF | 1.0 | 0.0 | 5.5 | 1.0 | -0.77e |
| D94S/G94E | CD | 1.0 | 1.0 | 5.0 | 0.0 | -0.75e |
|  | EF | 1.0 | 0.3 | 5.7 | 0.5 | -0.76e |
| $\Delta\alpha$PV | CD | 1.0 | 1.0 | 5.0 | 0.0 | -0.74e |
|  | EF | 1.0 | 0.0 | 5.0 | 1.0 | -0.77e |

$q_{eff}$ is used for the MSA calculations. The partial charges of different oxygen types are adapted from the Amber ff14SB force field. Specifically, the backbone oxygen charge is from SER O atom type, the hydroxyl oxygen charge is from SER OG atom type, the carbonyl oxygen charge is from GLU OD atom type and water oxygen charge is from the TIP3P water O atom type.

**Table S7:** Contact frequency of ion and CD site residues.

| System |  |  |  |  |  |  |
| --- | --- | --- | --- | --- | --- | --- |
| αPV:Ca<sup>2+</sup> | E62 100% | **E59 100%** | F57 100% | D53 100% | D51 100% | **S55 99%** |
| αPV:Mg<sup>2+</sup> | E62 100% | **E59 100%** | F57 100% | D53 100% | D51 100% | **S55 97%** |
| S55D/E59D αPV:Ca<sup>2+</sup> | E62 100% | F57 100% | **D55 100%** | D53 100% | D51 100% | |
| S55D/E59D αPV:Mg<sup>2+</sup> | E62 100% | F57 100% | **D55 100%** | D53 100% | D51 100% | |
| D94S/G98E αPV:Ca<sup>2+</sup> | E62 100% | E59 100% | F57 100% | D53 100% | D51 100% | S55 99% |
| D94S/G98E αPV:Mg<sup>2+</sup> | E62 100% | E59 100% | D53 100% | D51 100% | F57 99% | S55 98% |
| $\Delta\alpha$PV:Ca<sup>2+</sup> | E62 100% | E59 100% | F57 100% | D53 100% | D51 100% | S55 99% |
| $\Delta\alpha$PV:Mg<sup>2+</sup> | E62 100% | E59 100% | D53 100% | D51 100% | F57 99% | S55 98% |

Residues within 3Å of ions in the CD site are considered. Sorted from higher to the lower frequencies. Average of three sets of simulations are reported.

**Table S8:**
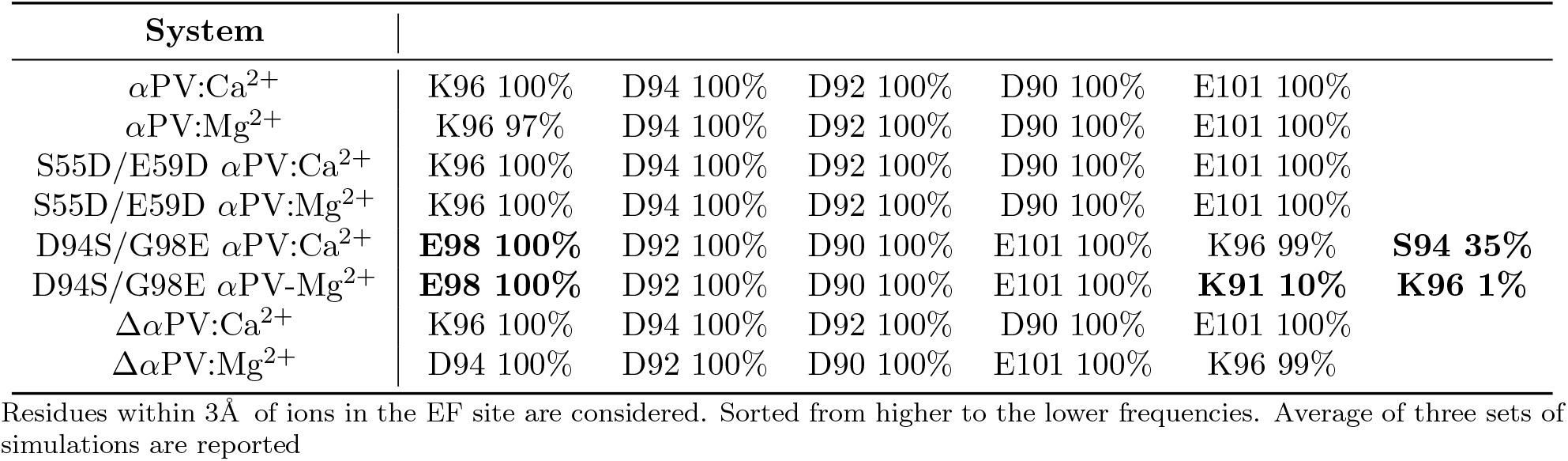
Contact frequency of ion and EF site residues.

### S8.3 Supplemental Methods

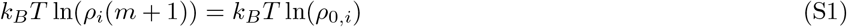

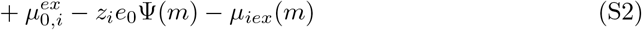

In Eq. S1, *ρ*_*i*_ is the density of the i-th ion type in the binding site, *ρ*_0,*i*_ is the density of the i-th ion in the bath (assumed to remain constant), 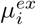 is the excess chemical potential of the i-th ion in the binding site, 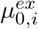 is the excess chemical potential of the i-th ion in the binding site (also constant),*z*_*i*_ is the integer charge of the i-th ion species, Ψ is the Donnan potential for the binding site, and 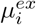 is the chemical potential of the i-th ion in the binding site.

### S8.4 Supplementary Results

*Protein backbone RMSDs* To assess the variability of other simulated systems with the experimental crystal structures and also to compare and contrast between different systems studied, we calculated the backbone RMSDs of protein. All the ion bound forms converged around 1Å (*≈*2Å for Δ*α*PV) in all the ion bound forms approximately after 100ns, where as the apo forms around 2Å after 600ns (*≈*5Å for Δ*α*PV). The backbone RMSDs shows that the presence of ions (Ca^2+^ or Mg^2+^) dampen the structural fluctuations of the variants compared to the apo forms (Fig. S1). While the difference between the apo and ion bound forms is statistically significant in all PV variants studied, no statistical difference between Ca^2+^ and Mg^2+^ bound forms was evident. Additionally, RMSD calculations suggest that the AB domain stabilizes the intact complex as its holo state RMSD is considerably smaller than the Δ*α*PV(*≈*5Å for truncated apo vs. the *≈*2Å for the other apo systems and *≈*2Å for truncated ion bound vs. *≈*1Å for other ion bound forms). Other pronounced structural variations are discussed further in sections 3.3. Overall, the structural similarities and converged RMSDs suggest that our simulation protocol can reproduce the conformational ensembles of ion-bound forms and also likely accurately model the apo-forms of the *α*PV variants for which no structural data are available.

*Water coordination of Ca*^2+^ We additionally assessed the number of waters coordinated by Ca^2+^ while bound to the CD and EF hand loops (Fig. 2E-H). We relate the cumulative number of protein and water oxygens in panels Fig. 2I-J to determine the total coordination number of Ca^2+^. In the CD hand, only the S55D/E59D *α*PV variant (Fig. 2G) reflects a bound water, which is roughly 2.2 Å from the Ca^2+^. In this capacity, the bound water in S55D/E59D *α*PV may help offset the loss of interaction of a bidentate residue E59 and coordinate D59 with the ion. In the EF site, all *α*PV structures except D94S/G98E *α*PV present one bound water in the coordination site Fig. 2H. The D94S/G98E *α*PV count is 0.5, signifying that a single water is bound for about one-half of the simulation time.

*Coordination shell in Mg*^2+^ *bound systems* In the Mg^2+^ bound systems we demonstrate through radial distribution of protein oxygens that the proteins bind tighter to the Mg^2+^ ion compared to the Ca^2+^. In either binding site, the first oxygen coordination shell is at *≈*1.7 Å for all the four systems (0.5 Å less than the Ca^2+^ bound systems) (Fig. 2E, F). This indicates that protein has to rearrange to interact with the ions (compared to Ca^2+^), which might come at a thermodynamics cost. The number of coordinating oxygens in this CD site for the S55D/E59D *α*PV is five, where as for all other systems it is six (Fig. 2G). In the EF site, five oxygens are coordinating with Mg^2+^ in all four systems (Fig. 2H). Overall, we show that, on average, two fewer oxygens are coordinating the Mg^2+^ (in both sites combined) compared to Ca^2+^ bound systems, which partly explains the lower affinity of Mg^2+^ to *α*PV systems compared to Ca^2+^. This is in line with the other earlier observations that the Mg^2+^ coordination is lower than Ca^2+^ in PVs [37]. Moreover, the water coordination in the Mg^2+^ bound systems is very similar to the Ca^2+^ bound ones, except that the first hydration shell is closer in the Mg^2+^ bound systems compared to the Ca^2+^ bound systems (*≈*0.5Å lower) (Fig. 2A-D).

For S55D/E59D *α*PV, the 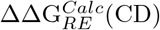 values attributed to the CD loop residues (L50-E62) summed to *≈ −*8 kcal/mol (see Table S2), suggesting that Ca^2+^ binding is more favorable in the S55D/E59D *α*PV site relative to the WT. This is not unexpected, given that the S55D substitution introduces an additional unit negative charge into the binding site. Meanwhile, 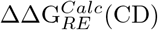 for D94S/G98E *α*PV is *≈*13 kcal/mol, which we found surprising given that the amino acid substitutions reside in the adjacent EF hand. Lastly, the score difference between the Δ*α*PV and WT holo states was approximately *≈*12 kcal/mol, which suggests Ca^2+^ binding was less favorable for the truncated species. Similarly, 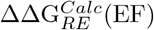 for the EF hand residues spanning D90 to F102 were *≈*14 kcal/mol for the S55D/E59D *α*PV suggesting that the Ca^2+^ binding in this site is unfavorable; where as for the D94S/G98E *α*PVit is *≈*38 kcal/mol suggesting the unfavorable binding of Ca^2+^, which trends correctly with the experimental data. Altogether, the 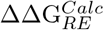 contributions arising from both CD and EF hands were *≈* 6 kcal/mol for S55D/E59D *α*PV and *≈* 51 kcal/mol for D94S/G98E *α*PV. Nevertheless, these scores from the CD and EF hands did not consistently rank the variants by affinity Table S2, which suggests that non-local contributions play a significant role in the binding scores.

*Specific hydrogen bonding interactions* The solvation differences between apo and holo states are driven by conformational changes upon ion binding. We therefore analyzed hydrogen bonds between amino acid side chains to determine if any may help stabilize the holo state. In Fig. S9 we report that for the hydrogen bonds that change between the apo and holo states, a greater number are formed during Ca^2+^ binding as opposed to breaking. This includes the formation of S55-E59 (CD site) and K96-D94 (EF site) hydrogen bonds in the holo forms and two hydrogen bonds that are lost upon ion binding in all systems; (S65-D76 that connects D-helix with the loop that is bridging D and E helices and S55-D51 that form part of the CD loop) (Fig. S9). However, in the S55D/E59D *α*PV interactions of S55 with E59 and D51 do not exist since the residues S55 and E59 were mutated to D55 and D59, respectively (Fig. S9). The D94S mutation in D94S/G98E *α*PV abolished the K96-D94 interaction, which we speculate results in high mobility of the EF site loop. Instead, a hydrogen bond between S94 and D90 is formed. In the Δ*α*PV system, the lack of AB negates the S78-D22 interaction.

*Glycine and glutamic acid play significant roles in tuning the dynamics of αPV helicity* In this study we observed that amino acid type and the location are very critical in *α*PVs. Proline and glycine are typical helix breakers [68, 50]. Our data show glycine significantly increase dynamics of *α*PV proteins as 4 out of 10 residues in the EF domain loop are glycine. For this reason the EF domain loop is highly flexible (glycines are the most flexible of all residues[50]) compared to the CD site loop in all *α*PVs, including the S55D/E59D *α*PV. The instability of the EF site loop due to the presence of extra glycine and aspartic acids is also reflected in the MM/GBSA reorganization energies As shown in Table S2, the 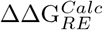 for the CD site loop residues is consistently more favorable than the EF site residues in all variants studied by at least 50 kcal/mol. Additionally, there are two glycines in the D-helix at positions 64 and 69. In particular, G64 is acting as a helix breaker in the D-helix as the helicity of residues from 65-70 is reduced by *≈*35% (Fig. 5C). The D-helix is part of the *α*PV hydrophobic core and interacts directly with the cation, therefore, cation binding in the CD site counter the effects of G64 and imparts stability to D-helix. Hence it is apparent that glycine increases the dynamics of *α*PV helicity and therefore may be a mechanism for increasing affinity.

### S8.5 Limitations

Limitations of our approach include the limited MD simulation length (microsecond), in addition to approximations associated with the MD and MM/GBSA methods (detailed in the review [49]). The time scale of binding, folding and unfolding etc. are usually on the order of milliseconds to seconds [85], compared to our microsecond-length simulations. The other significant limitation is that MM/GBSA is not sufficiently accurate to predict the solvation energies of ions, which is crucial to assess the absolute/relative binding affinities of the ions to *α*PV [49]. We attribute this to the inherent approximations in this end point method, such as the implicit treatment of solvent, neglect of entropy, and the sensitivity of the method to the solute dielectric constant and the challenges in accurately modeling the non-polar contribution to the solution free energy [86, 87, 88]. Force field deficiencies might also contribute to the inaccuracies in the binding affinity estimation, particularly those pertaining to describing interactions with divalent ions like Ca^2+^[89, 29].

Although, we used 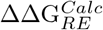 values from the MM/GBSA analysis to rank order all the variants according to their affinities, we omitted the apo state of Δ*α*PV for these comparisons. We believe Δ*α*PV was problematic because this simulation was started from the completely ordered holo structure by removing Ca^2+^, since the apo state crystal structure of the Δ*α*PV was unavailable; experimental studies report that the Δ*α*PV apo structure is completely disordered [18]. However, in our simulations we only observed partial unfolding of the structure, which warrants more simulation time that is beyond the scope of this study.

We also reported in Section 3.1 correlations between RMSF of the EF site loop residues and the affinity of ions for Ca^2+^. Although we could provide estimates for the enthalpy of the conformations comprising the apo and holo ensembles, the entropic contributions were inconclusive. We anticipate there are two significant contributions to entropy changes, including the apparent decrease in configurational entropy of the holo state relative to the apo state suggested by the RMSFs and the anticipated increase in solvent configuration entropy from desolvation of EF hand. It has been suggested that the increase in solvent configurational entropy likely predominates the loss of entropy from decreased conformational fluctuations [47].

Lastly, our MSA model indicates that the chemical potential is rather insensitive to the number of bound oxygens (8 is favored over 5 by less than 1kcal/mol). It is important to recognize that our MSA implementation does not account for quantum effects like polarization or orbital geometries that could favor certain coordination numbers and geometries [38, 54].

## Acronyms

*α*PV: *α*-parvalbumin. 1–20, 22, 23, 27–35
*β*PV: *β*-parvalbumin. 2, 3, 14, 27
Ca^2+^: calcium. 1–9, 11–33, 35
CaM: calmodulin. 16
CBP: Ca-binding protein. 1, 2, 16
CBPs: Ca-binding proteins. 3, 18
Cl^*−*^: chloride. 6
K^+^: potassium. 6
Mg^2+^: magnesium. 1–7, 9, 11–26, 28–33
MM/GBSA: molecular mechanics generalized Born approximation. 1, 3–7, 11, 13–16, 18–21
MSA: mean spherical approximation. 1–3, 6, 19, 21, 27
PV: Parvalbumin. 1, 2, 16, 19
RMSD: root mean squared deviations. 3
RMSF: root mean squared fluctuations. 3
TnC: troponin C. 3, 15, 16, 19
WT: wild-type. 3, 5, 6, 11, 13, 14, 16, 33

## Notes

### Competing Interest Statement

The authors have declared no competing interest.

